# Non-random interactions within and across guilds shape the potential to coexist in multi-trophic ecological communities

**DOI:** 10.1101/2021.11.29.470335

**Authors:** David García-Callejas, Oscar Godoy, Lisa Buche, María Hurtado, Jose B. Lanuza, Alfonso Allen-Perkins, Ignasi Bartomeus

**Affiliations:** Estación Biológica de Doñana (EBD-CSIC), Seville, Spain; Instituto Universitario de Ciencias del Mar (INMAR), Departamento de Biología, Universidad de Cádiz, E-11510, Puerto Real, Spain; Departamento de Ingeniería Eléctrica, Electrónica, Automática y Física Aplicada, ETSIDI, Technical University of Madrid, 28040 Madrid, Spain

## Abstract

Theory posits that the persistence of species in ecological communities is shaped by their interactions within and across trophic guilds. However, we lack empirical evaluations of how the structure, strength, and sign of biotic interactions drive the potential to coexist in diverse multi-trophic communities. Here we model community feasibility domains, a theoretically-informed measure of multi-species coexistence probability, from grassland communities comprising more than 45 species on average from three trophic guilds (plants, pollinators, and herbivores). Contrary to our hypothesis, increasing community complexity, measured either as the number of guilds or community richness, did not decrease community feasibility. Rather, we observed that high degrees of species self-regulation and niche partitioning allow maintaining larger levels of community feasibility and higher species persistence in more diverse communities. Our results show that biotic interactions within and across guilds are not random in nature and both structures significantly contribute to maintaining multi-trophic diversity.

## Introduction

Ecological communities are complex systems in which individuals of different species interact in a myriad of context-dependent ways, generating emergent properties that are not evident from the isolated study of their elements (Levin, 1998). Understanding these emergent properties, such as community stability or resilience (Meerbeek *et al.*, 2021) is key for strengthening the scientific basis of ecosystem conservation and restoration (Moreno-Mateos *et al.*, 2020). An important dimension of community stability is the potential for different species to coexist, i.e., to maintain viable populations in the same local community. However, obtaining a mechanistic understanding and quantifying the coexistence of multiple species in nature is a complex task because of the numerous processes that operate at the species and community levels.

Within a single trophic level, theoretical and empirical work on competitive interactions has shown that the degree of self-regulation relative to the strength of interspecific interactions is a key factor in shaping coexistence. Competitive communities are more stable when intraspecific interactions are high relative to interspecific ones (Buche *et al.*, 2022; Chesson, 2000; Levine & HilleRisLambers, 2009). The degree of overlap in resource use between species is also assumed to shape pairwise coexistence relationships, with implications for other emergent properties such as different ecosystem functions (Albert *et al.*, 2022; Godoy *et al.*, 2020). Likewise, structural properties, such as modularity for antagonistic networks, and nestedness for mutualistic ones, have been shown to promote community stability in bipartite communities (Rohr *et al.*, 2014; Stouffer & Bascompte, 2011; Thébault & Fontaine, 2010). However, all these insights have been derived from communities of single interaction types, either antagonistic or mutualistic. This progress contrasts with increasing evidence that different interaction types contribute synergistically to the emergent properties of ecological communities (Evans *et al.*, 2013; Losapio *et al.*, 2021; Simha *et al.*, 2022). A natural next step is, therefore, to study how coexistence is achieved in communities of increasing complexity, where interactions of different signs and strengths are intertwined across guilds. Disentangling this conundrum requires combining detailed empirical data with robust theoretical models.

Empirical studies documenting simultaneously multiple interaction types across several guilds are scarce. Early studies relied on binary networks that document the presence or absence of a given interaction (Bastolla *et al.*, 2009; Kéfi *et al.*, 2015). This approach can be refined by assigning interaction strengths inferred through indirect methods or expert opinion (e.g., Pocock *et al.* (2012); see García-Callejas *et al.* (2018) for a review). While these approaches are useful first approximations, documenting interactions quantitatively at finer scales, and over multiple communities, is essential for understanding the variability in community structure and dynamics (Banašek-Richter *et al.*, 2009), as recently shown in an agricultural context (Morrison *et al.*, 2020). Interactions within and across guilds have been mostly integrated into the context of mutualisms between plants and pollinators by incorporating competitive interactions to these bipartite networks (Bastolla *et al.*, 2009; Gracia-Lázaro *et al.*, 2018; Wang *et al.*, 2021), but the concept can be generalised to any kind of interaction (Godoy *et al.*, 2018; Seibold *et al.*, 2018). However, even when these intra-guild interactions are considered, an unrealistic solution has been to model them as constant across all species (i.e. a mean-field approach) (Bastolla *et al.*, 2009; Rohr *et al.*, 2014; Saavedra *et al.*, 2013). Although this approach is justified from a theoretical point of view because intra-guild interactions are notoriously difficult to quantify directly, this solution is suboptimal and lacks biological realism. There is widespread evidence that differences among species in phenological timing and resource use modulate variation in intra-guild interaction strengths (CaraDonna *et al.*, 2020; Morales-Castilla *et al.*, 2015). Consequently, such processes can generate differences in intra-guild network structures that, in turn, can influence their potential to maintain species diversity (Barabás *et al.*, 2016).

In parallel to empirical limitations in realistically quantifying diverse species interactions, mathematical tools for integrating this complexity in community-level frameworks are still under development (García-Callejas *et al.*, 2018; Godoy *et al.*, 2018; Pilosof *et al.*, 2017). Classic modelling approaches from single-interaction networks can be adapted to deal with multiple interactions (García-Callejas *et al.*, 2018; Mougi & Kondoh, 2012). However, they rely on adequate estimations not only of species interactions, but also of intrinsic growth rates, which are considerably difficult to quantify even for simplified communities (Bartomeus *et al.*, 2021), and potentially other parameters in more complex models (e.g., Gauzens *et al.* (2020)). Recent advances taking a structuralist approach provide an alternative to evaluating the role of species interactions in promoting or hindering multi-species coexistence (Godoy *et al.*, 2018; Saavedra *et al.*, 2017).

Conceptually, the coexistence of an arbitrary number of species in a community requires the existence of a steady state in which all species maintain positive abundances (i.e., the community is feasible) and return to it following perturbations (i.e., is stable *sensu* May (1972)). These conditions, while not equivalent, are tightly related, such that the existence of a feasible steady state implies its stability under a wide array of conditions (Barabás *et al.*, 2016; Gibbs *et al.*, 2018) given by the interplay between interaction strengths, their variability, and asymmetry (Bunin, 2017). The structuralist framework is built around the concept of feasibility and is often used to approximate coexistence in the absence of more detailed mechanistic insights or empirical information (Saavedra *et al.*, 2017). In this framework, the structure of species interactions, without other ancillary information, shapes the probabilistic opportunities to coexist for the different species in a given community by quantifying the so-called feasibility domain. The main prediction from the structuralist approach is precise: the larger the feasibility domain, the more likely the community can persist without any species going extinct. Indeed, communities with larger feasibility domains can withstand larger fluctuations in species vital rates without losing species (Song *et al.*, 2018). A key advantage of this probabilistic approach is that it further allows the derivation of probabilities of persistence (or its complement, exclusion) for individual species (Saavedra *et al.*, 2020). Thus, the structuralist approach emerges as a powerful tool to explicitly link estimations of persistence at the species and community levels, and to bridge theoretical studies on community stability and empirical quantifications of species interactions in diverse multi-trophic communities.

Here, we combine recent advances in the field of structural stability with unique field observations from nine Mediterranean grassland communities over two years involving a total of 108 taxa and their different types of interactions: plant-herbivore (antagonistic), plant-pollinator (mutualistic), and the intra- and interspecific competitive interactions within guilds (plants, pollinators, and herbivores). With this combination, we provide the first empirical exploration of how quantitative interaction structures within and across trophic guilds drive the persistence of biodiversity in natural conditions. Our first hypothesis is that the opportunities to coexist in our study system will be negatively related to the number of guilds considered and the overall richness of the community. In the absence of further mechanisms, a higher number of species entails higher degrees of niche overlap within and across guilds, which would imply weaker niche differences and fewer opportunities to coexist (Adler *et al.*, 2007; Buche *et al.*, 2022). This relationship, however, is expected to be modulated by the structure of species interactions in natural communities (Jacquet *et al.*, 2016). Therefore, our second hypothesis is that interaction structure within and across guilds will also shape the opportunities to coexist in multi-trophic communities. For testing these hypotheses, we define and analyse three different parameterisations of intra-guild interactions, and we compare the opportunities to coexist in the observed communities to randomised counterparts. Our third question asks whether network properties capture the variability in opportunities to coexist across the observed communities. In particular, we hypothesise that stronger self-regulation and a higher degree of resource partitioning across species will be related to higher opportunities to coexist in multi-trophic communities. We explicitly test these hypotheses both at the species and community levels of organisation.

## Methods

### Data collection

We conducted our study in a Mediterranean grassland community in Doñana National Park, SW Spain (37º 04’ 01.5”N, 6º 19’ 16.2” W). We set up 9 plots of 8.5 m^2^ across an area of 2680 ha (Fig. 1) from which we documented 1) direct competitive interactions among plants, 2) direct interactions between plants and pollinators, and 3) direct interactions between plants and herbivores. Because our study system is dominated by plants and insects that feed upon them (i.e. pollinators and herbivores), we expect these guilds to be the most relevant for the dynamics of the community, as the abundance of predators (e.g. spiders, mantis) or larger animals is relatively low. For plant-plant interactions, we obtained the number of local co-occurrences between plant individuals, sampling 36 focal individuals of each plant species per plot and their plant neighbours at a radius of 7.5 cm. This radius is a standard distance used in previous studies to measure competitive interactions among annual plant species (Levine & HilleRisLambers, 2009; Mayfield & Stouffer, 2017), and it has been validated to capture the outcome of competitive interactions at larger scales (1 m^2^) under locally homogeneous environmental conditions (Godoy & Levine, 2014). Interactions between plants and pollinators or herbivores were sampled from the emergence of the earliest flowers (February) to the decay of the latest ones (June) in 2019. During 2020, the length of the field season was the same, but we could not sample for five weeks in March/April 2020 due to COVID-19 restrictions. However, such differences in sampling effort did not seem to influence our results as trends were consistent across both years (see results).

**Fig. 1:**
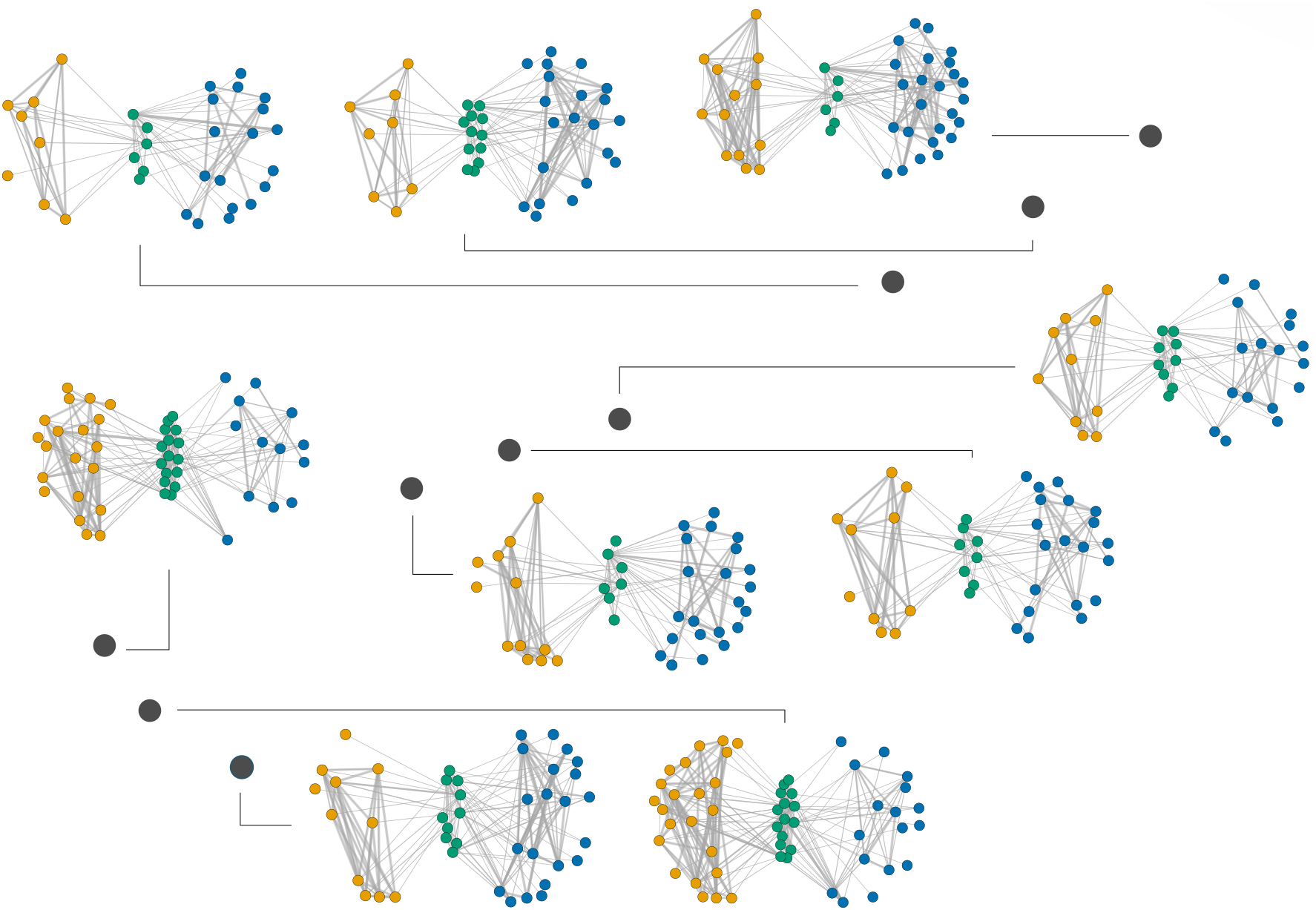
Approximate spatial configuration of the sampled networks. For reference, we show the 9 networks of 2019, with pollinators in orange (leftmost sets of nodes), plants in green (middle), and herbivores in blue (rightmost). Lines represent interactions observed or estimated from field information.

We recorded the number of floral visits to each plant species by sampling each plot for 30 min weekly for a total of 148.5 hours in 2019 and bi-weekly (therefore missing two sampling intervals) for a total of 54 hours in 2020. We only recorded floral visitors that contacted plant’s reproductive organs (stigma and/or anthers). Hence, we assume they are effective pollinators with positive population-level effects on plants, and refer to them that way throughout the text. Antagonistic interactions between plants and herbivores were sampled in parallel to the pollinator survey for 36 min on each plot, for a total of 76 hours in 2019 and 70 in 2020. Herbivores were annotated when observed on plant stems, leaves, or flowers.

From these field observations, we obtained 18 normalised block interaction matrices **A**_*n,t*_ (9 plots × 2 years). We assumed that these represent independent local communities given the spatial separation between plots (100 m on average), species turnover of the annual plants and the associated insect community, and the annual dynamics of the system. These matrices characterise the interaction structure of each local community, including intra- and inter-guild interactions, and are the inputs of the structural methods described below. The matrices are defined, for a given plot *n* and year *t*, as

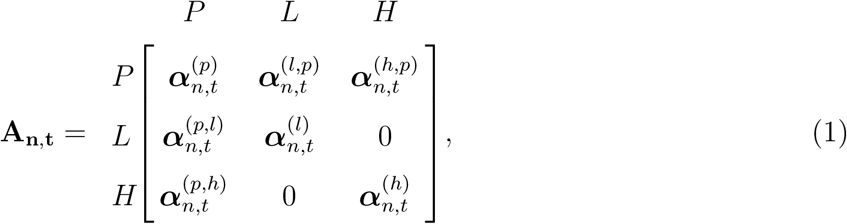

where *P* = plants, *L* = pollinators, and *H* = herbivores. The ***α*** elements represent the different sub-matrices (or blocks) of the community, e.g. ***α***^(*p*)^ represents the matrix of plant-plant interactions, ***α***^(*l,p*)^ the matrix of pollinator effects over plants, and so on. The different ***α*** submatrices have different sign structure depending on the interaction type: effects between plants and pollinators are positive in both directions 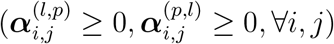, effects between plants and herbivores are antagonistic 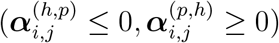, and intra-guild effects are negative 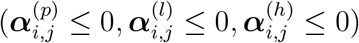.

### Multiple approaches to estimate community interactions

Estimating the occurrence and strength of interactions among the members of a guild is a pervasive problem in studies of ecological networks, in particular when individuals are highly mobile and/or difficult to track in the field (e.g., arthropod pollinators and herbivores). For these reasons, rather than providing a single characterisation, we present three different ones reflecting how the field of community ecology has evolved in the last decades. In our first parameterisation, named mean-field in the recent literature (Bastolla *et al.*, 2009; Rohr *et al.*, 2014; Saavedra *et al.*, 2013), we followed the implementation of previous works analysing competition mainly in plant-pollination networks (Bastolla *et al.*, 2009; Rohr *et al.*, 2014; Saavedra *et al.*, 2013). In this parameterisation, intra-guild competition affects all species equally and symmetrically (i.e., all competitive interaction coefficients are equal to a constant value). In our second parameterisation, we considered more structured intra-guild competition, which has recently been shown to modify the expected patterns of species persistence by generating mutualism-competition trade-offs (Gracia-Lázaro *et al.*, 2018; Wang *et al.*, 2021). We estimated the degree of asymmetric competition between plant species based on our spatially explicit field observations, and between pollinators (or herbivores) based on the feeding requirements of their larval stages and on their nesting requirements. This second parameterisation assumes that the structure of intra-guild competition relies on resource use. However, it overlooks phenological constraints commonly observed in natural communities (Olesen *et al.*, 2011), known to decouple interactions in time, decreasing net competition (Duchenne *et al.*, 2021). Therefore, in our third parameterisation we incorporated phenological overlap to the resource overlap axis of the second parameterisation. In the Supplementary Section “Interaction Matrices” we describe in detail the process of constructing these intra-guild matrices, of which we obtained one per plot and year for every guild and every parameterisation.

Interactions between individuals of different guilds (plants-herbivores and plants-pollinators) were obtained from the field observations described in the section “Data Collection”, and normalised to ensure comparable coefficients with the intra-guild matrices (Supplementary Section “Interaction Matrices”). For each plot and year, we thus obtained both intra- and inter-guild inter-action matrices for plants, herbivores, and pollinators, including three different parameterisations of intra-guild competition. For each parameterisation, this resulted in a total of 6 interaction matrices: three intra-guild communities ***α***^*p*^, ***α***^*l*^, ***α***^*h*^; two two-guild communities (one formed by plants and herbivores (***α***^*p*^, ***α***^*h*^, ***α***^*p,h*^, ***α***^*h,p*^), and one formed by plants and pollinators (***α***^*p*^, ***α***^*l*^, ***α***^*p,l*^, ***α***^*l,p*^)); and the overall community represented by the full block-matrix **A** (Eq. (1)). We further generated randomised counterparts of these different matrices to evaluate the effects of the observed interaction structure on our coexistence metrics (see Supplementary Section “Interaction Matrices” for details on the randomisation process).

### Community feasibility and species’ exclusion ratios

The potential for a given structure of species interactions to sustain a feasible community is quantified via its feasibility domain, whose relative volume ranges in the interval [0,0.5] (Song *et al.*, 2018). A large feasibility domain volume indicates a higher potential to accommodate variations in species growth rates while maintaining feasibility and vice-versa. Its mathematical definition is given in Song *et al.* (2018) and discussed in the Supplementary Section “Feasibility Metrics”; hereafter, we refer to this relative volume simply as “feasibility domain” for brevity. We calculated the feasibility domain of each of our interaction networks, i.e. for each community ***A***_*n,t*_ we calculated the feasibility domain of all sub-communities of one guild and two guilds, as well as of the full multi-trophic community. We further estimated the range of intrinsic growth rates contained within the feasibility domain of our communities, confirming that they agree with theoretical and ecological expectations (Fig. S5). The above calculations are based on the assumption that the dynamics of the communities studied can be reasonably well approximated with a linear Lotka-Volterra or equivalent models (Saavedra *et al.* (2017), but see Cenci & Saavedra (2018) for the extension of the structural approach to non-linear models).

Feasibility domains can potentially be highly asymmetric, meaning that some species are much closer to being excluded from a feasible community than others (Grilli *et al.*, 2017; Tabi *et al.*, 2020). However, analysing the feasibility domain alone does not provide information on the relative vulnerability of species to exclusion. To quantify these outcomes, we developed a novel structural measure for the likelihood of a species being excluded from a feasible community. This metric is a ratio that quantifies the relative probability of being the first species excluded, given the observed interaction matrix, compared to a situation in which all species have the same probability of being excluded. These probabilities are obtained by quantifying the proportion of the feasibility domain’s volume that is closer to the border where only a given species *i* goes excluded (see Supplementary Section “Feasibility Metrics” for the mathematical definition and an extended discussion). This novel metric, the *species exclusion ratio*, is bounded between (0, ∞), and is a good proxy of the overall likelihood of exclusion of a species in the absence of further information on, e.g.. intrinsic growth rates.

### Structural metrics for species and communities

We hypothesise that, in diverse systems, species with weaker self-regulation and higher niche overlap will generally be less likely to coexist (Adler *et al.*, 2007; Barabás *et al.*, 2016; Buche *et al.*, 2022; Godoy *et al.*, 2017), and this will be reflected in the feasibility domain of the community. We calculated two complementary structural network metrics for the species and community levels to test these tenets in multi-trophic communities. First, we quantified the degree of self-regulation of a species as its diagonal dominance. Diagonal dominance is a matrix property that is satisfied when diagonal elements are larger than the sum of non-diagonal elements. Here we used a continuous version, i.e. the difference between the diagonal and the sum of non-diagonal elements. This species-level metric is averaged for obtaining the average degree of diagonal dominance in a community, *d*:

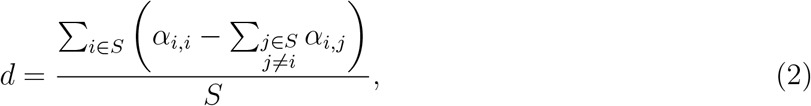

where *S* is the number of species in the community represented by the interaction matrix *α*. Similarly, we obtained species-level overlap and its community average, assuming that the degree of overlap in pairwise interactions is an appropriate proxy of niche overlap. To avoid confusion, we refer hereafter to interaction overlap. Species-level interaction overlap is itself an aggregated property, derived from the overlap between each pair of species, calculated using the Morisita-Horn dissimilarity index (Horn, 1966) implemented in the R package `vegan` v2.6-2 (Oksanen *et al.*, 2022). We posit that the net overlap of species *i* in a community, *o_i_*, is best represented by the sum of pairwise overlaps with every other species:

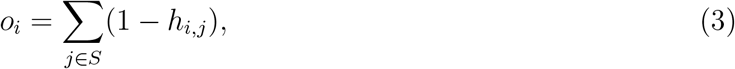

where *h* is a Morisita-Horn dissimilarity matrix obtained from a given interaction matrix. From Eq. 3, we obtained separately the interaction overlap of each species i for its intra-guild competition matrices and its inter-guild interaction matrices. The community-level metric is, again, the average of species-level overlaps in the community:

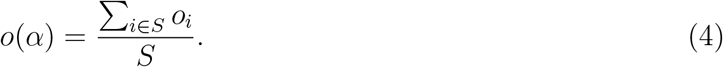

### Statistical analyses

We analysed the relationship between coexistence outcomes and structural metrics using regression models. For testing whether increased complexity decreased opportunities to coexist (first hypothesis), we analysed the relationship between the feasibility domain (for the community-level analyses) or species exclusion ratios (for the species-level analyses) and the number of trophic guilds accounted for using a Type III Analysis of Variance. We used linear models to explore the relationship between the feasibility domain (or species exclusion ratios) and species richness.

In addition, by explicitly considering the three different types of intra-guild competition matrices, we also explored our second hypothesis that interaction structure influences the opportunities to coexist in our system. Furthermore, we compared the observed feasibility domains (or species exclusion ratios) to the distributions given by the randomised communities. To analyse the relationship between our coexistence outcomes and structural metrics (our third hypothesis), we analysed the full communities, including plants, herbivores, and pollinators, with the most structured intra-guild interactions (resource use and phenological overlap). Using this data, we implemented linear mixed models with feasibility domain as response variable (log-transformed, to account for its non-negativity), and community-level diagonal dominance and interaction overlap (differentiating intra-guild and inter-guild overlap) as independent variables, taking the plot identity as a spatial random factor. For the species-level analyses, we similarly took species exclusion ratios (log-transformed) as response, and diagonal dominance, intra-guild overlap, and inter-guild overlap as independent variables. In this model, we also added species guild as a covariate and again included plot identity as a random factor. We implemented these models using the `lmerTest` package v3.1-3 in R (Kuznetsova *et al.*, 2017). We scaled all numeric variables and checked model fits with the tests provided in the package `DHARMa` v0.4.5 (Hartig, 2021).

## Results

In our two years of sampling, we documented 214 unique interactions among plants, 110 between plants and pollinators, and 160 between plants and herbivores. In this period, we observed interactions between 108 taxa, of which 17 were plants, 46 herbivores, and 45 pollinators. Of these, 53 taxa representing 49% of the records were identified at the species level (17 plants, 16 pollinators, and 20 herbivores), and 51 % as morphospecies (Table S1). The included taxa span diverse life-history strategies, such as grasses (e.g. *Hordeum marinum*) and forbs (e.g. *Leontodon maroccanus*) in the annual plant guild, solitary bees (e.g. from genera *Andrena, Lasioglossum*), flies (e.g. genera *Sphaerophoria, Musca*), or Lepidoptera (e.g. genera *Lasiocampa, Thymelicus*) within the pollinator guild, and sap feeders (e.g. Hemiptera from genera *Aelia* or *Aphis*), pollen feeders (e.g. Coleoptera from genera *Malachius* or *Psilothrix*) or leaf-eaters (e.g. Gastropoda from genera *Theba* or *Cochlicella*) within the herbivore guild (Table S1).

The frequency distribution of the 270 unique interactions observed across trophic guilds was highly skewed; e.g., 128 interactions were observed less than five times. Plant richness was positively correlated with that of pollinators across communities (Spearman’s *ρ* = 0.78, S = 214, p-value < 0.01) but not with herbivores (Spearman’s *ρ* = −0.23, S = 1125, p-value = 0.35). Richness values averaged 48 taxa and ranged from 35 to 57 in the least and most diverse communities, respectively. The sampling coverage of all interaction types, quantified via rarefaction curves, was > 80% in all local communities and > 90% in most of them (Supplementary Section “Rarefaction Analyses”).

Contrary to our first hypothesis that the higher the complexity of the local communities, the lower the opportunities for species to coexist, we found that neither of the complexity proxies influenced feasibility domains (Fig. 2). In particular, we found no significant differences in the feasibility domain of communities considering one, two, or three guilds (Type III Analysis of Variance, number of guilds: F = 1.54, df = 2, p-value = 0.22). Likewise, community richness did not significantly affect our communities’ feasibility domain (Table S2). However, at the species level, the degree of complexity influenced the exclusion ratios, but in the opposite direction to our expectation. Specifically, both the number of guilds in a community (Fig. S2 and Table S3) and community richness (Fig. S3 and Table S4) showed a statistically significant negative relationship with species’ exclusion ratios, alongside significant interaction effects between richness, types of intra-guild competition, and species guild. Overall, these results suggest that, on average, species are comparatively more likely to persist in more diverse communities.

**Fig. 2:**
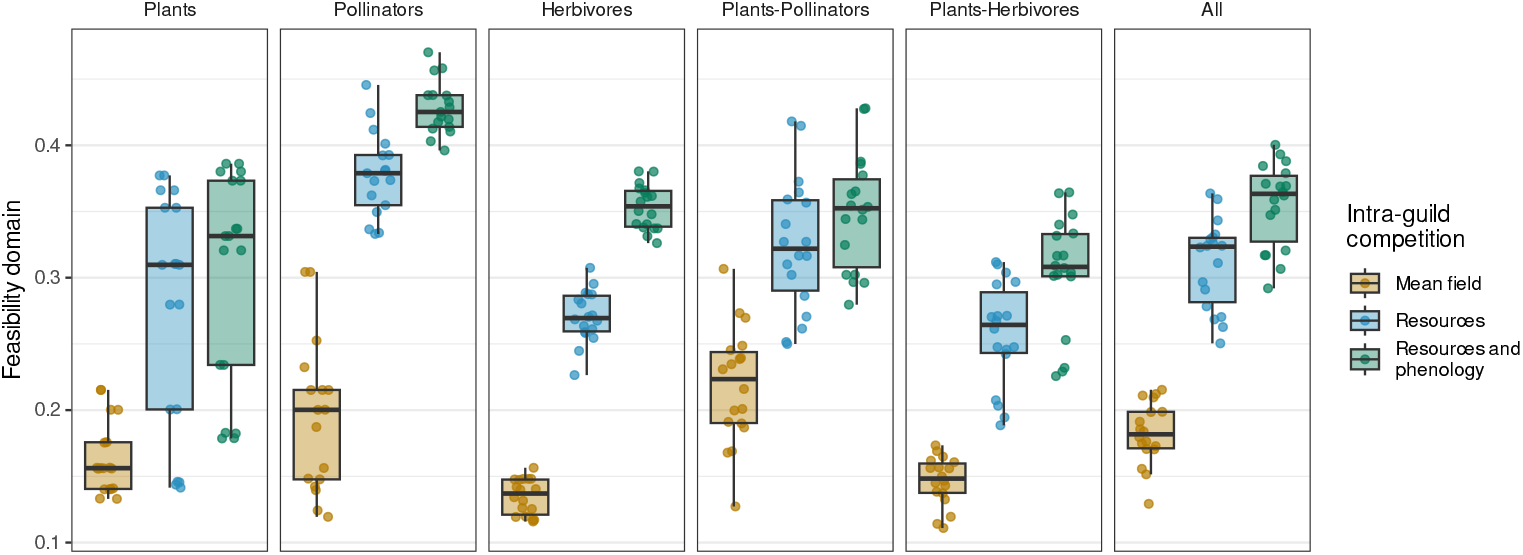
Feasibility domain volumes of each community or subcommunity, for the three different parameterisations of intra-guild competition matrices (feasibility domain volumes range in the interval [0,0.5]). In the boxplots, the horizontal black line represents the median, the lower and upper hinges correspond to the 25th and 75th percentiles, and the vertical lines extend to the largest/smallest value up to 1.5 times the interquartile range (distance between 25th and 75th percentiles). N = 18 communities in each boxplot (9 plots × 2 years).

Regarding our second hypothesis, we found that different parameterisations of the intra-guild competition matrices resulted in significant differences in feasibility domains (Fig. 2; Type III Analysis of Variance, intra-guild type: F = 176.85, df = 2, p-value < 0.001; Table S2). In particular, feasibility domains were lowest for communities with mean-field competition and highest for those with resource competition mediated by phenological overlap. While the mean-field parameterisation generated the lowest feasibility domains in all situations, community rankings based on their feasibility domain were reasonably well maintained across parameterisations (Fig.S1; Spearman’s *ρ* = 0.57, S = 416, p-value = 0.015). Therefore, although the mean-field approach underestimates the potential for coexistence compared with other parameterisations, it is still useful for characterising relative differences across communities. At the species level, exclusion ratios were not different on average across guilds, with plants, pollinators, and herbivores displaying similar distributions. The type of intra-guild competition significantly influenced the variability of species’ exclusion ratios rather than the mean (Fig. 3, Fig. S2). Mean-field competition communities displayed much higher variability in this metric than the other two parameterisations. Overall, these results suggest that the feasibility domains of communities with mean-field competition are smaller due to higher variability in species’ exclusion ratios, such that a small subset of species with high exclusion ratios (the upper hinges of the boxplots in Fig. 3) drag the feasibility domains of these communities down.

**Fig. 3:**
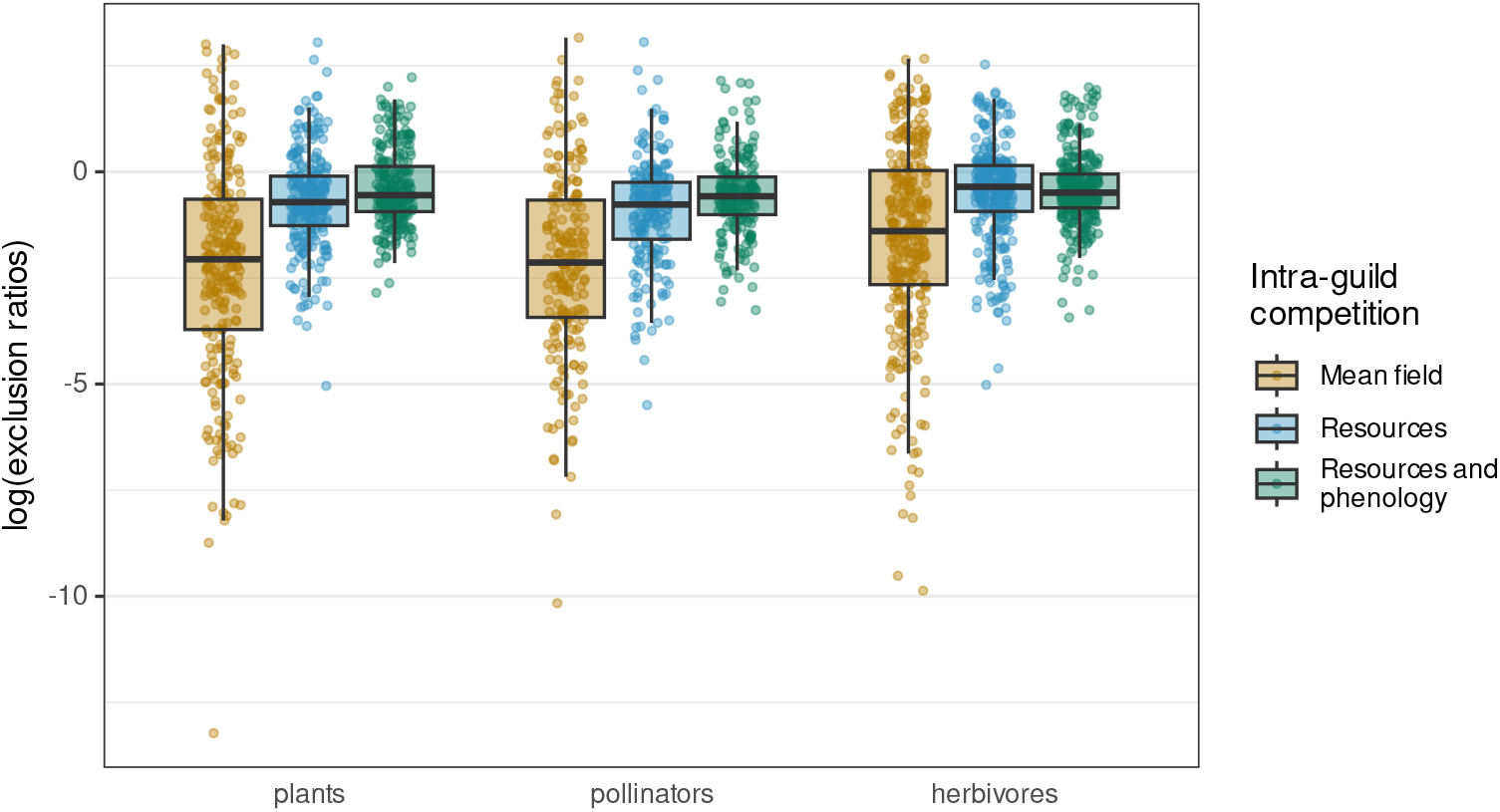
Species’ exclusion ratios (log-transformed) for the different guilds and the three intra-guild parameterisations. Here we show the results for the full communities, considering the three guilds. In all parameterisations, N = 204 for plants, 201 for pollinators, and 303 for herbivores.

In further agreement with our second hypothesis, the whole architecture of multi-trophic systems, including intra- and inter-guild species interactions, influenced the opportunities to coexist. We found that randomising interaction structures resulted for every parameterisation in smaller feasibility domains compared to the structures observed in the field. Considering the full communities with three trophic guilds, only 4 communities out of 54 (18 per parameterisation) fell within the 95% interval of the null distributions (Fig. 4). Similarly, we found that such randomisations also increased the average and the variability of species’ exclusion ratios in all situations (Fig. S4, Table S5).

**Fig. 4:**
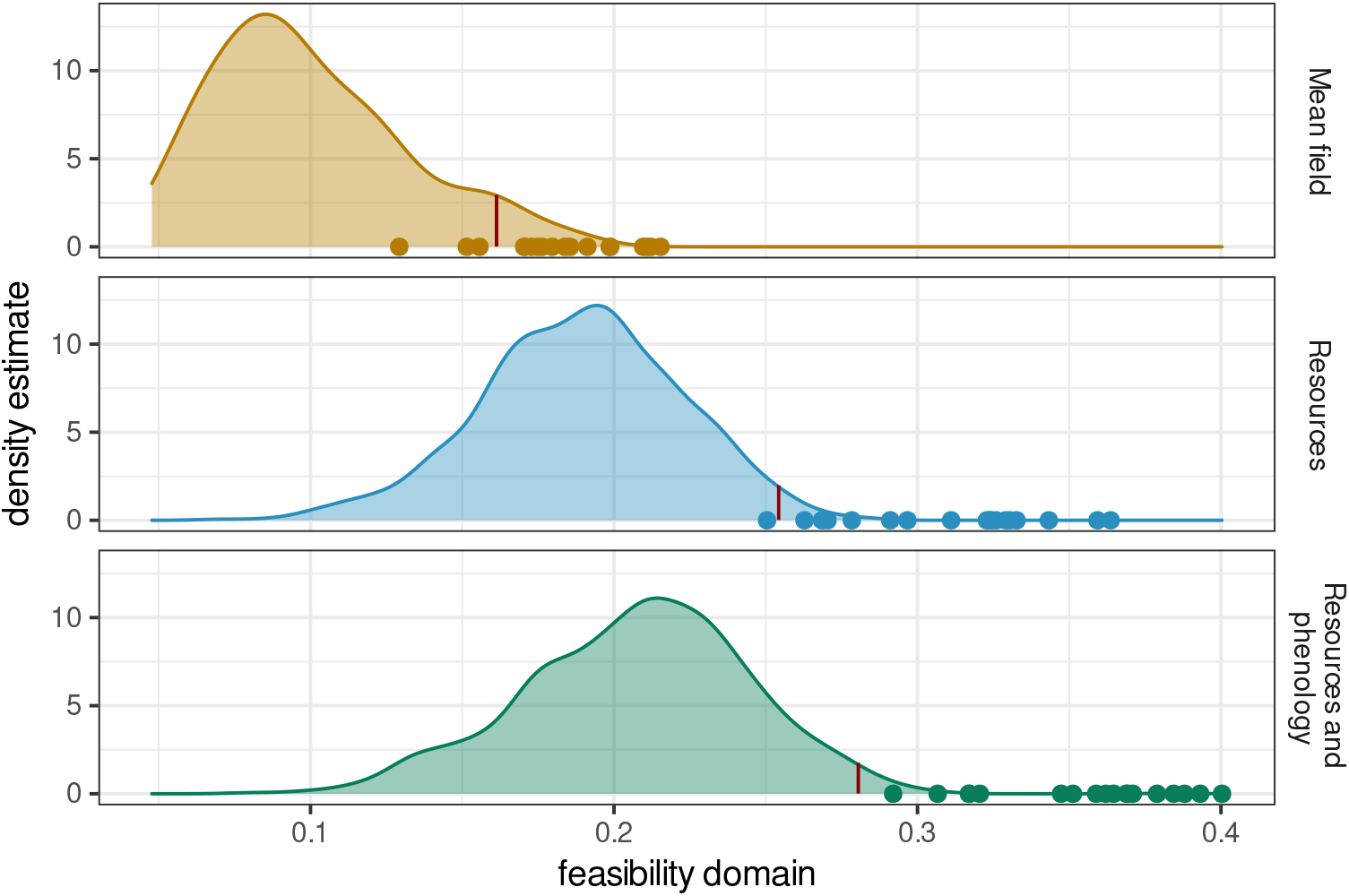
Feasibility domain of the observed communities (N=18 in each panel), and the distribution of feasibility domain values from the randomised communities. Red vertical lines represent the 97.5% percentile of the null distributions. For reference, the observations here correspond to the right-most panel in Fig. 2, i.e., to the full communities, including plants, pollinators, and herbivores.

Finally, and supporting our third hypothesis, we found that two of the three proposed network properties were related to the feasibility domain in our full communities. Specifically, we observed a positive relationship between the average degree of diagonal dominance and the feasibility domain, a negative relationship with the average degree of intra-guild interaction overlap, and no statistically significant relationship with inter-guild interaction overlap (Table 1, Fig. 5). The effect size of the two significant metrics was qualitatively similar (Table 1). Likewise, species exclusion ratios showed qualitatively similar trends with species-level metrics (Table S6). For all trophic guilds (plants, pollinators, and herbivores), diagonal dominance significantly negatively affected species exclusion ratio. In contrast, intra-guild and inter-guild interaction overlap had significant positive effects. The three metrics had qualitatively comparable effect sizes.

**Table 1:** Coefficients of the Linear Mixed Model relating feasibility domain (log-transformed) with average diagonal dominance, intra- and inter-guild interaction overlap. The three independent variables were not correlated (all Variance Inflation Factors < 1.07). The estimated *σ_plot_* is 0.023. N = 18.

|  | estimate | std.error | t | p.value |
| --- | --- | --- | --- | --- |
| Intercept | -1.095 | 0.05 | -21.875 | $< 0.001$ |
| avg_diagonal_dominance | 0.208 | 0.055 | 3.757 | 0.002 |
| avg_intraguild_overlap | -0.198 | 0.074 | -2.659 | 0.03 |
| avg_interguild_overlap | -0.011 | 0.055 | -0.206 | 0.839 |

## Discussion

Our results provide evidence, using real-world communities, for two fundamental and tightly related questions in community ecology. First, the opportunities to coexist do not decrease with increasing community richness or a higher number of trophic guilds in our study system. Second, the structure of interactions between species of the same guild and between species of different guilds are both key for maintaining the opportunities to coexist, especially via niche partitioning and densitydependent mechanisms of self-regulation (Barabás *et al.*, 2017). These insights rest upon field observations from highly diverse communities comprising three distinct trophic guilds: annual plants, their pollinators, and their insect and gastropod herbivores.

**Fig. 5:**
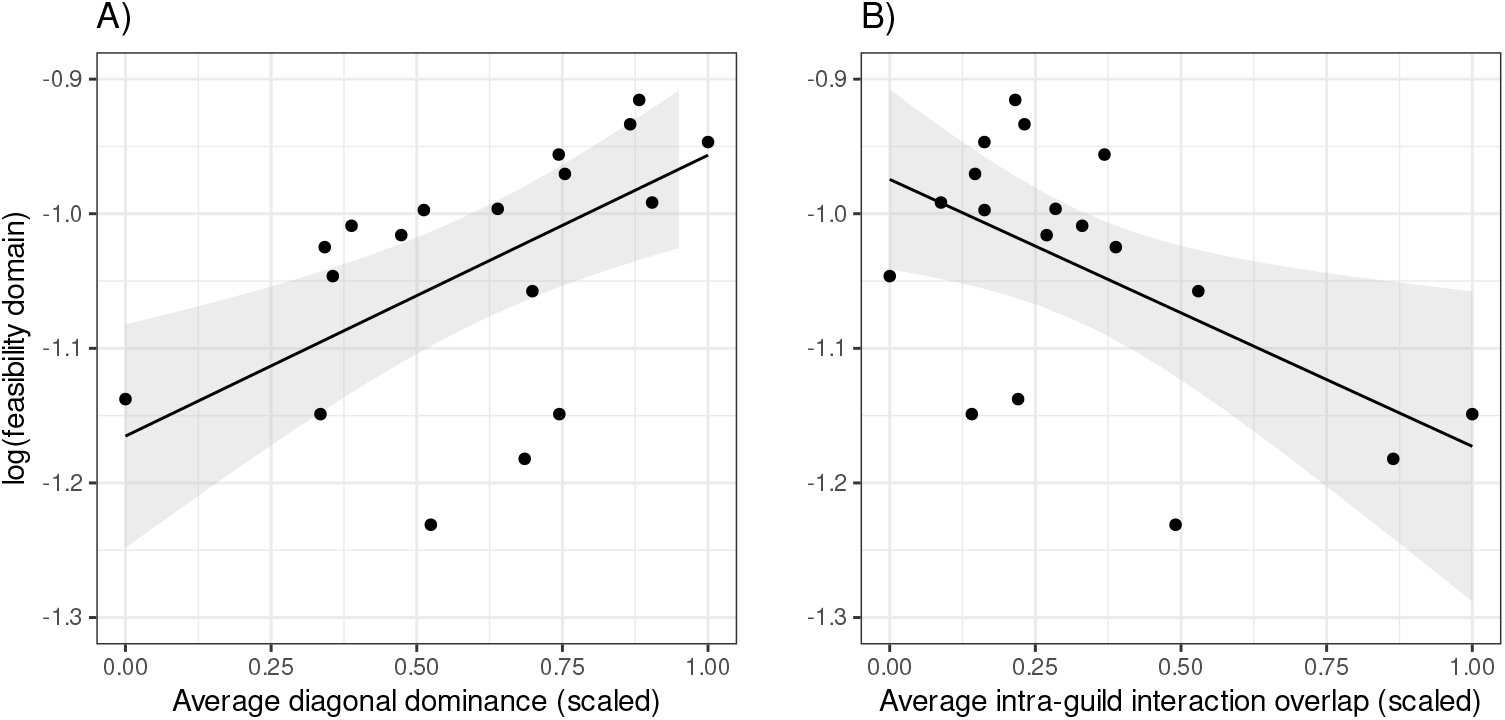
Relationship between the feasibility domains of our full communities (log-transformed, N=18 in each panel), average diagonal dominance (panel A), and average intra-guild interaction overlap (panel B). The observations here correspond to the full communities with intra-guild competition matrices parameterised by resource use and phenological overlap, i.e. the right-most boxplot in Fig. 2 and the lowest panel in Fig. 4.

The number of guilds or species considered in our communities does not influence the potential to maintain multi-species coexistence (Fig. 2). Theoretical work has demonstrated that the chances of finding stable (May, 1972) or feasible (Dougoud *et al.*, 2018) communities with random interaction structures decrease with increasing richness. The key to reconciling the theoretical expectations with our observations is that this theory applies to *random interaction structures*. Our results show that the ecological mechanisms structuring our communities out of randomness apply to different levels of organisation, from single guilds (Fig. 2) to multi-trophic communities (Fig. 4). These insights are, to our knowledge, the first empirical demonstration of the role of inter-action structures at different levels in shaping multi-trophic coexistence. Coexistence is, however, a multidimensional concept, exceedingly difficult to test in the field (Clark *et al.*, 2019), and thus here we maintain a probabilistic approximation. In our study system, as in most studies on empirical communities, we cannot easily measure the different dimensions of coexistence, in particular asymptotic stability, without strong assumptions on species’ intrinsic growth rates. Therefore, it is essential to keep in mind that our results refer to the probabilistic potential for feasibility and coexistence, expressed by the volume of the feasibility domains of our communities.

The hypothesis that realistic interaction structures enhance different facets of community stability with respect to random configurations has been repeatedly brought up to explain the apparent persistence of empirical communities (Jacquet *et al.*, 2016; Medeiros *et al.*, 2020). However, recent studies show conflicting results on whether community feasibility in particular is more likely under realistic structural constraints (Dougoud *et al.*, 2018; Grilli *et al.*, 2017; Saavedra *et al.*, 2016; Serván *et al.*, 2018). For example, Grilli *et al.* (2017) showed that random food webs have larger feasibility domains than empirical ones, whereas mutualistic networks do not show this difference. Dougoud *et al.* (2018), on the other hand, argued that only realistic interaction structures confer feasibility to species-rich food webs. The disparities in these works stem partly from considering different systems (food webs, competitive or mutualistic communities, either empirical or random), null models and assumptions, and configurations of interaction strengths. Our study provides a way to integrate these contrasting outcomes by linking species and community feasibility to structural constraints with a clear ecological interpretation (Fig. 5). For example, if species in an observed community show stronger interaction overlap than expected, we would expect a smaller feasibility domain in the observed community than in their random counterparts. This could happen for example if interactions are sampled in periods of resource pulses, or where certain resources are temporarily more available, as may be the case in the winter predator-prey network of the Białowieza forest analysed by Saavedra *et al.* (2016).

To complement the community insights, we developed parallel analyses at the species level. The exclusion ratio is a novel probabilistic metric that quantifies how likely a species is to be the first excluded from its community in the face of a perturbation, relative to a situation where all species are equally likely to be excluded. The exclusion ratio, importantly, does not rely on numerical simulations of steady states (as in, e.g., Saavedra *et al.* (2020)) and, therefore, can be reliably estimated given only the interaction matrix of a local community. In an applied context, for communities whose dynamics can be well approximated by Lotka-Volterra or equivalent frameworks, the species exclusion ratio may therefore help identify locally vulnerable species or guilds, as well as hint at community-level outcomes. Indeed, the combined analysis of species and community metrics facilitates the interpretation of our results: in our more structured intra-guild parameterisations, lower interaction overlap and higher self-regulation lead to smaller and relatively homogeneous species exclusion ratios (Fig. 3), which entail comparatively larger feasibility domains in the community (Fig. 4). Thus, the relative homogeneity of species exclusion ratios emerges as a key metric for understanding and comparing feasibility across communities. We hypothesise that higher variability in exclusion ratios is more likely to be found in communities in which species properties lead to higher asymmetry in species interactions’ strengths. Such asymmetry usually is observed in systems with comparatively larger variability in body sizes (Atkins *et al.*, 2015), trophic guilds, and life history strategies (Germain *et al.*, 2016), as opposed to study systems like ours.

Our study lacks estimations of intrinsic growth rates for every taxon in the community. This information would allow predicting tangible outcomes of which species can maintain positive populations rather than estimating probabilistic opportunities to coexist. However, obtaining such fine-scale estimations is logistically unfeasible for field observations of diverse communities. Given these limitations and the equally stringent data requirements of more mechanistic population dynamics models (e.g., Gauzens *et al.* (2020); Valdovinos (2019)), we reinforce the idea that estimating feasibility domains, alongside the potential range of growth rates compatible with feasibility (Fig. S5), can be a useful probabilistic approximation to multi-species coexistence (Saavedra *et al.*, 2020). This information, importantly, can be obtained directly from community interaction matrices when assuming that populations follow Lotka-Volterra or equivalent dynamics (Saavedra *et al.*, 2017), potentially incorporating non-linear functional forms (Cenci & Saavedra, 2018). Further-more, pairwise interactions across guilds can be quantified from interaction frequencies if these are assumed to be a good proxy of overall species effects, which is generally the case for insect pollinators (Vázquez *et al.*, 2012). In the case of intra-guild interactions, proxies of competition can lead to pairwise interaction effects based on different dimensions of resource overlap (Morales-Castilla *et al.*, 2015). Using interaction frequencies as a proxy for overall interaction strengths is nevertheless a first approximation in the absence of better-resolved data (Novella-Fernandez *et al.*, 2019). For example, further refinements of this methodology can account for varying per-capita efficiencies in pollen transportation for pollinators or in plant damage for herbivores. Incorporating these and other mechanisms, through their effects on community interactions, will allow better disentangling the relative importance of different ecological processes underlying the feasibility of natural communities.

Overall, our study highlights the need to adopt an integrative view of ecological communities because the structure of biotic interactions both within and across guilds is critical to shaping multi-species coexistence. By advancing in this integration, we identified the degree of niche partitioning and self-regulation within guilds as critical determinants of local species persistence and the feasibility of multi-trophic communities. We provide a fully operational framework to quantify these properties (degree of niche partitioning and self-regulation) from interaction matrices of any combination of interaction types, thus opening the door to compare on common grounds the potential to coexist across different community types. Across ecological communities, inter-action structures might vary due to additional factors, biotic or abiotic (e.g. invasive species, N deposition). Nevertheless, we show that for a highly diverse Mediterranean grassland, ecological interactions are structured in such a way as to maintain the opportunities for species to coexist across guilds and increasing complexity.

## Acknowledgements

We thank the Radical Community Ecology group(https://github.com/RadicalCommEcol/) for the fruitful discussions. DGC, IB, OG, and AA-P were funded by the Spanish Ministry of Science and Innovation (MICINN) and the European Social fund through the MeDiNaS (RTI2018-098888-A-I00), TASTE (PID2021-127607OB-I00), and ChaSisCOMA (PID2021-122711NB-C21) projects. Additionally, OG acknowledges financial support provided by the Spanish Ministry of Economy and Competitiveness (MINECO) and by the European Social Fund through the Ramón y Cajal Program (RYC2017-23666). MH acknowledges financial support provided by the Spanish Ministry of Science and Innovation through the FPI grant program (PRE 2019-088280).

## Data and code availability

The data and code used to generate the results of this study are available at https://doi.org/10.5281/zenodo.7527011.

