## Supplementary Material for "Non-random interactions within and across guilds shape the potential to coexist in multi-trophic ecological communities"

This document contains Supplementary Tables S1-S6, Supplementary Figures S1-S10, and Supplementary Sections "Interaction Matrices", "Feasibility Metrics", and "Rarefaction Analyses".

Table S1: Taxa included and number of interactions observed, summed across plots and years

| ID | guild | order | family | genus | species | interactions |
|---|---|---|---|---|---|---|
| BEMA | plants | Caryophyllales | Amaranthaceae | *Beta* | *macrocarpa* | 214 |
| CETE | plants | Gentianales | Gentianaceae | *Centaurium* | *tenuiflorum* | 1236 |
| CHFU | plants | Asterales | Asteraceae | *Chamaemelum* | *fuscatum* | 1235 |
| CHMI | plants | Asterales | Asteraceae | *Chamaemelum* | *mixtum* | 48 |
| HOMA | plants | Poales | Poaceae | *Hordeum* | *marinum* | 8093 |
| LEMA | plants | Asterales | Asteraceae | *Leontodon* | *maroccanus* | 8656 |
| MESU | plants | Fabales | Fabaceae | *Melilotus* | *sulcatus* | 752 |
| PAIN | plants | Poales | Poaceae | *Parapholis* | *incurva* | 1006 |
| PLCO | plants | Lamiales | Plantaginaceae | *Plantago* | *coronopus* | 46 |
| POMA | plants | Poales | Poaceae | *Polypogon* | *maritimus* | 499 |
| POMO | plants | Poales | Poaceae | *Polypogon* | *monspeliensis* | 282 |
| PUPA | plants | Asterales | Asteraceae | *Pulicaria* | *paludosa* | 677 |
| RAPE | plants | Ranunculales | Ranunculaceae | *Ranunculus* | *peltatus* | 18 |
| SASO | plants | Caryophyllales | Amaranthaceae | *Salsola* | *soda* | 12 |
| SCLA | plants | Asterales | Asteraceae | *Scorzonera* | *laciniata* | 62 |
| SOAS | plants | Asterales | Asteraceae | *Sonchus* | *asper* | 241 |
| SPRU | plants | Caryophyllales | Caryophyllaceae | *Spergularia* | *rubra* | 69 |
| Anastoechus_spp | pollinators | Diptera | Bombyliidae | *Anastoechus* | | 43 |
| Andrena_argentata | pollinators | Hymenoptera | Andrenidae | *Andrena* | *argentata* | 3 |
| Andrena_cinerea | pollinators | Hymenoptera | Andrenidae | *Andrena* | *cinerea* | 56 |
| Andrena_humilis | pollinators | Hymenoptera | Andrenidae | *Andrena* | *humilis* | 70 |
| Andrena_spp | pollinators | Hymenoptera | Andrenidae | *Andrena* | | 54 |
| Bombyliidae | pollinators | Diptera | Bombyliidae | | | 51 |
| Bombylius_major | pollinators | Diptera | Bombyliidae | *Bombylius* | *major* | 13 |
| Braconidae | pollinators | Hymenoptera | Braconidae | | | 6 |
| Calliphoridae | pollinators | Diptera | Calliphoridae | | | 16 |
| Chrysididae | pollinators | Hymenoptera | Chrysididae | | | 1 |
| Chrysotoxum_spp | pollinators | Diptera | Syrphidae | *Chrysotoxum* | | 4 |
| Colias_croceus | pollinators | Lepidoptera | Pieridae | *Colias* | *croceus* | 6 |
| Coscinia_spp | pollinators | Lepidoptera | Erebidae | *Coscinia* | | 6 |
| Culicidae | pollinators | Diptera | Culicidae | | | 1 |
| Cylindromyia_spp | pollinators | Diptera | Tachinidae | *Cylindromyia* | | 8 |
| Dilophus_spp | pollinators | Diptera | Bibionidae | *Dilophus* | | 500 |
| Diplazon_spp | pollinators | Hymenoptera | Ichneumonidae | *Diplazon* | | 1 |
| Empis_spp | pollinators | Diptera | Empididae | *Empis* | | 2 |
| Empis_tesellata | pollinators | Diptera | Empididae | *Empis* | *tesellata* | 21 |
| Episyrphus_balteatus | pollinators | Diptera | Syrphidae | *Episyrphus* | *balteatus* | 42 |
| Eristalis_spp | pollinators | Diptera | Syrphidae | *Eristalis* | | 20 |
| Eucera_spp | pollinators | Hymenoptera | Apidae | *Eucera* | | 58 |
| Euchloe_crameri | pollinators | Lepidoptera | Pieridae | *Euchloe* | *crameri* | 1 |

| | | | | | | |
|---|---|---|---|---|---|---|
| Eupeodes_corollae | pollinators | Diptera | Syrphidae | *Eupeodes* | *corollae* | 76 |
| Geometridae | pollinators | Lepidoptera | Geometridae | | | 2 |
| Lasiocampa_trifolii | pollinators | Lepidoptera | Lasiocampidae | *Lasiocampa* | *trifolii* | 58 |
| Lasioglossum_immunitum | pollinators | Hymenoptera | Halictidae | *Lasioglosum* | *immunitum* | 10 |
| Lasioglossum_malachurum | pollinators | Hymenoptera | Halictidae | *Lasioglossum* | *malachurum* | 104 |
| Lasioglossum_spp | pollinators | Hymenoptera | Halictidae | *Lasioglossum* | | 11 |
| Lomatia_spp | pollinators | Diptera | Syrphidae | *Lomatia* | | 32 |
| Melanopangonius_spp | pollinators | Diptera | Tabanidae | *Pangonius* | | 1 |
| Melanostoma_spp | pollinators | Diptera | Syrphidae | *Melanostoma* | | 2 |
| Musca_spp | pollinators | Diptera | Muscidae | *Musca* | | 166 |
| Nemotelus_spp | pollinators | Diptera | Stratiomyidae | *Nemotelus* | | 40 |
| Nephrotoma_spp | pollinators | Diptera | Tipulidae | *Nephrotoma* | | 2 |
| Odontomyia_spp | pollinators | Diptera | Stratiomyidae | *Odontomyia* | | 21 |
| Osmia_ligurica | pollinators | Hymenoptera | Megachilidae | *Osmia* | *ligurica* | 14 |
| Pangonius_spp | pollinators | Diptera | Tabanidae | *Pangonius* | | 16 |
| Pieris_brassicae | pollinators | Lepidoptera | Pieridae | *Pieris* | *brassicae* | 3 |
| Sarcophaga_spp | pollinators | Diptera | Sarcophagidae | *Sarcophaga* | | 21 |
| Scaeva_spp | pollinators | Diptera | Syrphidae | *Scaeva* | | 23 |
| Sphaerophoria_scripta | pollinators | Diptera | Syrphidae | *Sphaerophoria* | *scripta* | 180 |
| Sphaerophoria_spp | pollinators | Diptera | Syrphidae | *Sphaerophoria* | | 3 |
| Thymelicus_spp | pollinators | Lepidoptera | Hesperiidae | *Thymelicus* | | 10 |
| Ulidiidae | pollinators | Diptera | Ulidiidae | | | 216 |
| Vanessa_cardui | pollinators | Lepidoptera | Nymphalidae | *Vanessa* | *cardui* | 8 |
| Acrididae | herbivores | Orthoptera | Acrididae | | | 60 |
| Aelia_spp | herbivores | Hemiptera | Pentatomidae | *Aelia* | | 172 |
| Agriotes_spp | herbivores | Coleoptera | Elateridae | *Agriotes* | | 77 |
| Aiolopus_strepens | herbivores | Orthoptera | Acrididae | *Aiolopus* | *strepens* | 116 |
| Aleyrodidae | herbivores | Hemiptera | Aleyrodidae | | | 4 |
| Anthaxia_semicuprea | herbivores | Coleoptera | Buprestidae | *Anthaxia* | *semicuprea* | 4 |
| Aphis_fabae | herbivores | Hemiptera | Aphididae | *Aphis* | *fabae* | 348 |
| Brassicogethes_spp | herbivores | Coleoptera | Nitidulidae | *Brassicogethes* | | 3076 |
| Bruchidae | herbivores | Coleoptera | Bruchidae | | | 5 |
| Cantharis_coronata | herbivores | Coleoptera | Cantharidae | *Cantharis* | *coronata* | 70 |
| Cantharis_spp | herbivores | Coleoptera | Cantharidae | *Cantharis* | | 2 |
| Cassida_spp | herbivores | Coleoptera | Chrysomelidae | *Cassida* | | 4 |
| Centrocoris_spp | herbivores | Hemiptera | Coreidae | *Centrocoris* | | 1 |
| Cercopoidea | herbivores | Hemiptera | Cercopoidea | | | 2 |
| Cicadidae | herbivores | Hemiptera | Cicadidae | | | 1 |
| Cleta_ramosaria_caterpillar | herbivores | Lepidoptera | Geometridae | *Cleta* | *ramosaria* | 3 |
| Cochlicella_barbara | herbivores | Gastropoda | Geomitridae | *Cochlicella* | *barbara* | 7074 |
| Conocephalus_dorsalis | herbivores | Orthoptera | Tettigoniidae | *Conocephalus* | *dorsalis* | 36 |
| Cryptocephalus_spp | herbivores | Coleoptera | Chrysomelidae | *Cryptocephalus* | | 32 |
| Dociostaurus_jagoi | herbivores | Orthoptera | Acrididae | *Dociostaurus* | *jagoi* | 1 |

| | | | | | | |
|---|---|---|---|---|---|---|
| Dolycoris_spp | herbivores | Hemiptera | Pentatomidae | *Dolycoris* | | 1 |
| Geometridae_caterpillar | herbivores | Lepidoptera | Geometridae | | | 6 |
| Gryllus_bimaculatus | herbivores | Orthoptera | Gryllidae | *Gryllus* | *bimaculatus* | 14 |
| Lagorina_sericea | herbivores | Coleoptera | Meloidae | *Lagorina* | *sericea* | 18 |
| Lasiocampa_trifolii_caterpillar | herbivores | Lepidoptera | Lasiocampidae | *Lasiocampa* | *trifolii* | 820 |
| Lepidoptera_caterpillar | herbivores | Lepidoptera | | | | 96 |
| Malachius_bipustulatus | herbivores | Coleoptera | Melyridae | *Malachius* | *bipustulatus* | 102 |
| Meligethes_spp | herbivores | Coleoptera | Nitidulidae | *Meligethes* | | 114 |
| Meloidae | herbivores | Coleoptera | Meloidae | | | 1 |
| Miridae | herbivores | Hemiptera | Miridae | | | 38 |
| Mirini_spp | herbivores | Hemiptera | Miridae | *Mirini* | | 16 |
| Mordellidae | herbivores | Coleoptera | Mordellidae | | | 66 |
| Nymphalidae_caterpillar | herbivores | Lepidoptera | Nymphalidae | | | 2 |
| Oedemeridae | herbivores | Coleoptera | Oedemeridae | | | 112 |
| Otala_lactea | herbivores | Gastropoda | Helicidae | *Otala* | *lactea* | 10 |
| Oxycarenus_hyalinipennis | herbivores | Hemiptera | Lygaeidae | *Oxycarenus* | *hyalinipennis* | 4 |
| Phaedon_spp | herbivores | Coleoptera | Chrysomelidae | *Phaedon* | | 1 |
| Philaenus_spumarius | herbivores | Hemiptera | Aphrophoridae | *Philaenus* | *spumarius* | 2 |
| Psilothrix_viridicoerulea | herbivores | Coleoptera | Melyridae | *Psilothrix* | *viridicoerulea* | 2946 |
| Pyrrhocoris_apterus | herbivores | Hemiptera | Pyrrhocoridae | *Pyrrhocoris* | *apterus* | 1 |
| Scyphophorus_spp | herbivores | Coleoptera | Curculionidae | *Scyphophorus* | | 20 |
| Tessellana_tessellata | herbivores | Orthoptera | Tettigoniidae | *Tessellana* | *tessellata* | 1236 |
| Tetranychus_urticae | herbivores | Trombidiformes | Tetranychidae | *Tetranychus* | *urticae* | 1 |
| Theba_pisana | herbivores | Gastropoda | Helicidae | *Theba* | *pisana* | 4410 |
| Thysanoptera | herbivores | Thysanoptera | | | | 19 |

Table S2: Linear model for the effects of community richness (scaled in the range 0-1) and type of intra-guild competition on the log-transformed feasibility domain of the studied communities. For this model we considered all sub-communities of one or two guilds, as well as the full communities (N = 324). Model fit was checked with the R package `DHARMA` v0.4.5 (Hartig, 2021).

| term | estimate | std.error | t | p.value |
|---|---|---|---|---|
| Intercept | -1.72799 | 0.04438 | -38.93329 | < 0.001 |
| richness | -0.12757 | 0.09188 | -1.38837 | 0.16600 |
| intraguild.typeResources | 0.60037 | 0.06277 | 9.56502 | < 0.001 |
| intraguild.typeResources and phenology | 0.73618 | 0.06277 | 11.72868 | < 0.001 |
| richness:intraguild.typeResources | -0.09198 | 0.12994 | -0.70790 | 0.47953 |
| richness:intraguild.typeResources and phenology | -0.06292 | 0.12994 | -0.48423 | 0.62856 |

Table S3: Analysis of Variance (type III) for species' exclusion ratios as function of three categorical factors and their interactions: number of guilds (guild.num label), intra-guild parameterisation (intraguild.type), and guild of the species (sp.guild), as well as their interactions. N = 7458.

| term | sum of squares | Df | F | p-value |
|---|---|---|---|---|
| Intercept | 0.743 | 1.000 | 0.451 | 0.502 |
| guild.num | 669.772 | 2.000 | 203.335 | < 0.001 |
| intraguild.type | 6.548 | 2.000 | 1.988 | 0.137 |
| sp.guild | 5.892 | 2.000 | 1.789 | 0.167 |
| guild.num:intraguild.type | 359.802 | 4.000 | 54.616 | < 0.001 |
| guild.num:sp.guild | 149.388 | 4.000 | 22.676 | < 0.001 |
| intraguild.type:sp.guild | 18.330 | 4.000 | 2.782 | 0.025 |
| guild.num:intraguild.type:sp.guild | 105.434 | 8.000 | 8.002 | < 0.001 |
| Residuals | 12238.617 | 7431.000 | | |

Table S4: Linear model for species' exclusion ratios (log-transformed) as a function of community richness (scaled in the range 0-1), intra-guild parameterisation (intraguild.type label), and guild of the species (sp.guild), as well as their interactions. N = 7458. Model fit was checked with the R package `DHARMA` v0.4.5 (Hartig, 2021), and showed significant deviation in the Kolmogorov-Smirnov test for normality of residuals. However, we decided to keep this analysis as the visual inspection of the residuals displayed no apparent pervasive bias.

| term | estimate | std.error | t | p.value |
|---|---|---|---|---|
| Intercept | 0.351 | 0.092 | 3.814 | $< 0.001$ |
| richness.scaled | -3.595 | 0.166 | -21.670 | $< 0.001$ |
| intraguild.typeResources | -0.574 | 0.130 | -4.418 | $< 0.001$ |
| intraguild.typeResources and phenology | -0.671 | 0.130 | -5.160 | $< 0.001$ |
| sp.guildherbivores | 0.083 | 0.138 | 0.603 | 0.546 |
| sp.guildfloral visitors | 0.042 | 0.137 | 0.305 | 0.760 |
| richness.scaled:intraguild.typeResources | 3.104 | 0.235 | 13.230 | $< 0.001$ |
| richness.scaled:intraguild.typeResources and phenology | 3.623 | 0.235 | 15.443 | $< 0.001$ |
| richness.scaled:sp.guildherbivores | 0.721 | 0.244 | 2.957 | 0.003 |
| richness.scaled:sp.guildfloral visitors | 0.751 | 0.241 | 3.120 | 0.002 |
| intraguild.typeResources:sp.guildherbivores | 0.450 | 0.195 | 2.303 | 0.021 |
| intraguild.typeResources and phenology:sp.guildherbivores | 0.471 | 0.195 | 2.413 | 0.016 |
| intraguild.typeResources:sp.guildfloral visitors | 0.402 | 0.194 | 2.074 | 0.038 |
| intraguild.typeResources and phenology:sp.guildfloral visitors | 0.366 | 0.194 | 1.886 | 0.059 |
| richness.scaled:intraguild.typeResources:sp.guildherbivores | -1.406 | 0.345 | -4.079 | $< 0.001$ |
| richness.scaled:intraguild.typeResources and phenology:sp.guildherbivores | -1.677 | 0.345 | -4.865 | $< 0.001$ |
| richness.scaled:intraguild.typeResources:sp.guildfloral visitors | -1.547 | 0.340 | -4.548 | $< 0.001$ |
| richness.scaled:intraguild.typeResources and phenology:sp.guildfloral visitors | -1.572 | 0.340 | -4.621 | $< 0.001$ |

Table S5: Summary statistics (mean, standard deviation, minimum and maximum values) of the distributions of exclusion ratios (e.r. in the table) for the observed and randomised communities, differentiating across guilds and intra-guild parameterisations. These distributions are obtained from the full communities.

| guild | intra-guild interactions | type | mean e.r. | sd e.r. | min e.r. | max e.r. |
|---|---|---|---|---|---|---|
| pollinators | Mean field | null | 8.32579 | 43.13503 | 0.00000 | 514.43056 |
| pollinators | Mean field | observed | 1.03675 | 1.96776 | 0.00002 | 23.57585 |
| pollinators | Resources | null | 16.20503 | 69.88239 | 0.00000 | 789.12712 |
| pollinators | Resources | observed | 0.93185 | 1.18537 | 0.00411 | 21.25431 |
| pollinators | Resources and phenology | null | 14.65369 | 62.62634 | 0.00000 | 725.54739 |
| pollinators | Resources and phenology | observed | 0.94833 | 0.88812 | 0.03843 | 8.56900 |
| herbivores | Mean field | null | 10.21271 | 46.56078 | 0.00000 | 450.00000 |
| herbivores | Mean field | observed | 1.05625 | 1.83193 | 0.00002 | 14.34667 |
| herbivores | Resources | null | 24.02730 | 76.63680 | 0.00000 | 778.79569 |
| herbivores | Resources | observed | 1.05005 | 1.04650 | 0.00325 | 12.52897 |
| herbivores | Resources and phenology | null | 24.19445 | 78.12533 | 0.00000 | 836.00000 |
| herbivores | Resources and phenology | observed | 1.01198 | 0.95144 | 0.03227 | 7.32789 |
| plants | Mean field | null | 7.43829 | 43.00815 | 0.00000 | 452.24545 |
| plants | Mean field | observed | 0.92499 | 2.04661 | 0.00000 | 21.10259 |
| plants | Resources | null | 17.89564 | 67.40658 | 0.00000 | 688.13314 |
| plants | Resources | observed | 0.99535 | 1.42795 | 0.00647 | 21.21409 |
| plants | Resources and phenology | null | 18.85959 | 67.62882 | 0.00000 | 899.75954 |
| plants | Resources and phenology | observed | 1.02066 | 0.98941 | 0.02812 | 9.25334 |

Table S6: Linear mixed model for the effects of species-level metrics (scaled) on log-transformed exclusion ratios. The estimated $\sigma_{plot}$ is 0.1862. For this model we considered the full communities with intra-guild competition parameterised using resource use mediated by phenological overlap (i.e. the right-most boxplot of Fig. 2. N = 748). All covariates display Variance Inflation Factors (adjusted by degrees of freedom) < 2. Model fit was checked with the R package `DHARMA` v0.4.5 (Hartig, 2021), and showed significant deviation in the Kolmogorov-Smirnov test for normality of residuals. However, we decided to keep this analysis as the visual inspection of the residuals displayed no apparent pervasive bias.

| term | estimate | std.error | t | p-value |
|---|---|---|---|---|
| Intercept | -0.508 | 0.235 | -2.160 | 0.031 |
| sp.guildpollinators | 0.349 | 0.151 | 2.317 | 0.021 |
| sp.guildherbivores | 0.118 | 0.135 | 0.873 | 0.383 |
| diagonal_dominance | -1.208 | 0.282 | -4.284 | < 0.001 |
| total_intraguild_overlap | 1.540 | 0.290 | 5.310 | < 0.001 |
| total_interguild_overlap | 1.036 | 0.147 | 7.058 | < 0.001 |

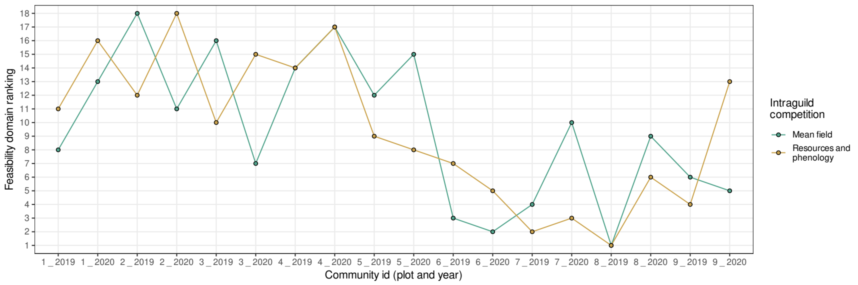


Fig. S1: Local communities ranked by their feasibility domain values in the two extreme parameterisations of intra-guild competition: mean field and resource use mediated by phenological overlap. The community with the highest rank in y-axis has the largest feasibility domain, and so on. Observations are the 18 local communities, i.e. one community per plot and year.

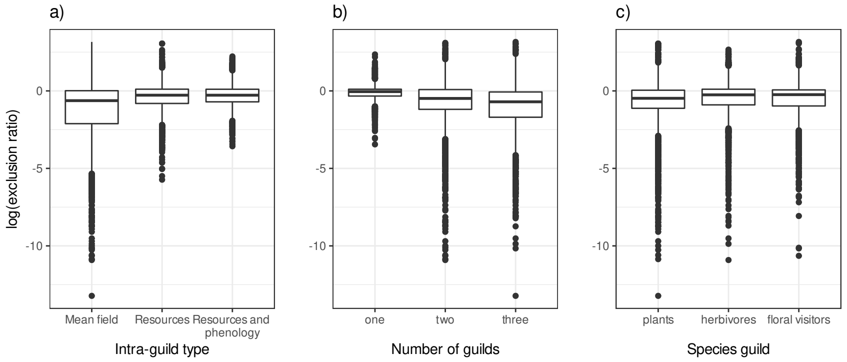


Fig. S2: Species exclusion ratios according to different factors: type of intra-guild parameterisation (panel a), number of guilds in the community (panel b), and guild of the species (panel c). N = 7458 in each panel. Boxplot components as in Fig. 2. See table S3 for the associated Analysis of Variance.

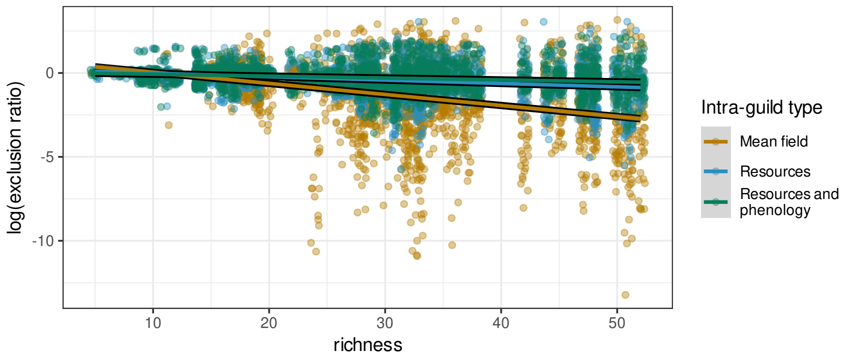


Fig. S3: Species exclusion ratios in relationship to community richness, for each intra-guild parameterisation. N = 7458. See table S4 for the associated model coefficients.

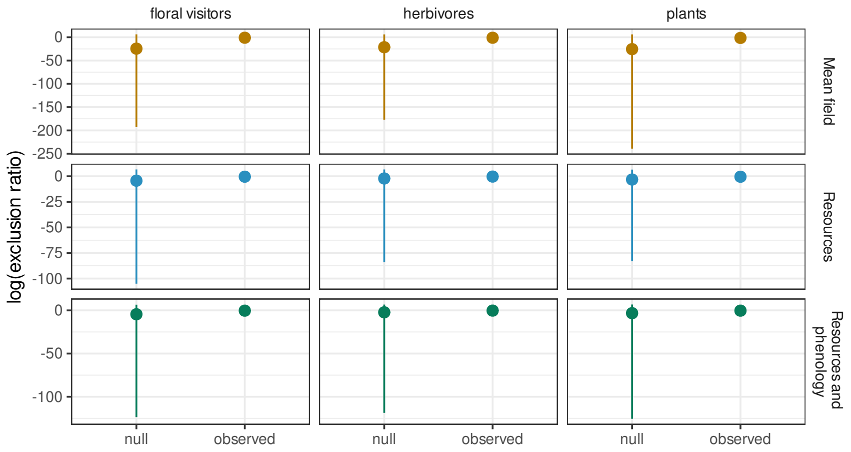


Fig. S4: Exclusion ratios summaries for observed and randomised (null) interaction structures. Shown are mean values (points) and maximum values (limit of the vertical lines). We display distributions this way because of the high peaks and long tails of the distributions, which difficult their full visualisation. Summary statistics can be consulted in Table S5.

### Supplementary Section: Interaction Matrices

Here, we describe in detail the three parameterisations of intra-guild competition matrices that we developed: Mean-field competition, resource-based competition, and resource-based competition mediated by phenology. Furthermore, we also discuss the integration of inter-guild interactions, and the network randomisation process.

##### Mean-field matrices

The most parsimonious set of matrices, those assuming mean-field, were derived following previous studies (Bastolla *et al.*, 2009; Saavedra *et al.*, 2013), by assigning a constant value $c_{inter} = 0.2$ to all inter-specific interactions between species, and a larger constant value $c_{intra} = 1$ to all intra-specific interactions, reflecting the assumption that individuals of a given taxon tend to compete more strongly with their conspecifics than with heterospecifics.

##### Resource-use matrices

Resource-use matrices are based on resource axes independent from those defined by plant-animal interactions because competition for plant resources among adult animals is already captured indirectly in inter-guild interactions. Therefore, we defined an axis of intra-guild competition based on larval and nesting requirements for animal species, and plant spatial associations between plant individuals as a proxy for competition for nutrients and water. For animals, we built two intermediate matrices, one for larval feeding requirements and another for nesting requirements. When two taxa share a larval feeding resource (nesting requirement) their inter-specific interaction coefficient was set to $c_{inter}$, otherwise inter-specific coefficients were zero; intra-specific interactions were always set to $c_{intra}$. The final matrix was obtained by averaging the values of these two matrices. Therefore, a given pair of species will display maximal inter-specific competition if they share both larval feeding requirements and nesting requirements. For plants, we simply standardised between 0 and 1 the number of observed spatial associations at 7.5 cm between species pairs, taking these as a proxy of inter-specific competition (Levine & HilleRisLambers, 2009; Mayfield & Stouffer, 2017). Intra-specific competition was assumed to be equal to $c_{intra}$, as in the mean-field situation. This was done to 1) maintain the same assumption in the animal and plant matrices, 2) because inter-specific values larger than intra-specific ones can give rise to numerical issues with the estimation of feasibility and structural metrics.

**Resource-use and phenological overlap matrices**

In this parameterisation we added phenological overlap to our resource-use matrices, as a common modulating factor for all intra-guild interactions. Note that in this case we assume that there are no preempting processes, i.e. the amount of resources such as soil, water, or food available to a species is not altered by earlier taxa. We calculated the phenological overlap between species $i$ and species $j$ in a given year as the activity period in which both species are observed relative to the full activity period of species $i$. We define the activity period of species $i$ in a given year as the interval between species $i$'s first and last observations recorded in the entire study area that year. Note that this implies that the phenological overlap between any two species can potentially be asymmetric. For weighing the resource-use matrix by phenological overlap, we multiplied element-wise both matrices. This assumes that the overlap in nesting requirements and larval feeding requirements of our species is correlated with the activity periods observed in the field.

In the last step common to all parameterisations, to standardise the resulting matrices we divided each element by the sum of all elements in the matrix, i.e. for plants:

$$\boldsymbol{\alpha}_{n,t}^{(p)} = \frac{\mathbf{b}_{n,t}^{(p)}}{\sum_{i,j}(\mathbf{b}_{n,t})_{ij}}, \tag{S1}$$

where $\mathbf{b}$ is the raw matrix, prior to normalisation. This normalisation implies that the elements of the resulting $\boldsymbol{\alpha}$ matrices sum to 1. In ecological terms, this means that every matrix has an identical weight in the full community $\mathbf{A}_{n,t}$, or in other words, every interaction has an identical net effect on the community. Also note that, up to this point, all matrices are non-negative. Thus, negative signs were imposed on the resulting $\boldsymbol{\alpha}$ matrices when appropriate: all intra-guild competition matrices, and herbivore effects on plants (see below). Although we acknowledge that intra-guild interactions may be positive in particluar occasions, here we considered only competition to simplify our analyses.

##### Inter-guild interactions

Given the observational data with the number of visits from a certain species of pollinator (or herbivore) $i$ to a certain plant species $j$, we obtained the normalised interaction intensity by dividing each element of the original bipartite matrix by the sum of all matrix elements (Eq. (S1)). As with the intra-guild matrices, this ensures inter-guild coefficients in the range [0,1] and that the sum of all elements in each matrix equals 1. Likewise, the sign structure (e.g. (+,+) relationships for plants-pollinators, (-,+) for plants-herbivores) is imposed directly over the resulting matrices. Overall, aside from the assumption of equal interaction effect on the full community, this assumes that the population-level interaction effect between a plant species and a pollinator or herbivore is given by the number of visits observed in the field (Vázquez *et al.*, 2005). This assumption is a sensible approximation given that all inter-guild interactions recorded are plant-arthropod interactions whereby all animals have body sizes $<$ 5cm, and that pollinator and herbivore interactions vary only in one order of magnitude (e.g. maximum number of interactions observed for herbivores is 7074 for *Cochlicella barbara* and for pollinators is 500 (*Dilophus* spp). Thus, the elements of the inter-guild matrices (e.g. $\boldsymbol{\alpha}_{n,t}^{p,h}$ for effects of herbivores over plants in plot $n$ and year $t$) represent population-level effects between any two taxa, a formulation that has already been used in the structuralist framework (Song *et al.*, 2018a).

##### Null matrices

To evaluate the effects of the observed interaction structure on multi-species coexistence, we generated randomised counterparts of each community and parameterisation. The null model used randomised interaction coefficients within any given block, thus conserving the overall connectance of the networks but randomising degree distributions and any other topological pattern. In these null communities, we did not consider forbidden interactions, which implies that a non-zero coefficient can potentially be assigned to an ecologically unfeasible interaction. We generated 100 null replicates of each interaction matrix, for a total of 2 years $\times$ 9 plots $\times$ 6 interaction matrices $\times$ 3 intra-guild parameterisations $\times$ 100 replicates = 32400 randomised communities.

### Supplementary Section: Feasibility Metrics

In this section we provide the mathematical definition of the structural stability metrics used in our study. First, the potential for a given structure of species interactions to sustain a feasible community is quantified via its feasibility domain, whose relative volume ranges in the interval [0,0.5] (Song *et al.*, 2018b). A large feasibility domain indicates a higher potential to accommodate variations in species growth rates while maintaining feasibility, and vice-versa. In mathematical terms, given an interaction matrix $\mathbf{A}$ for $S$ species, its feasibility domain is defined as

$$D_F(\mathbf{A}) = \{\mathbf{r} = N_1^* \boldsymbol{\nu_1} + ... + N_S^* \boldsymbol{\nu_s}, \text{with } N_1^* > 0, ..., N_S^* > 0\}, \tag{S2}$$

where the vector $\boldsymbol{\nu_j}$ is the negative of the $j$th columns of the interaction matrix $\mathbf{A}$, the vector $\mathbf{r}$ represents species' intrinsic growth rates in a classic Lotka-Volterra system, and $\mathbf{N}^* = -\mathbf{A}^{-1}\mathbf{r}$ is the vector of abundances when the feasible equilibrium exists (Song *et al.*, 2018b). This quantity is then normalised with respect to the full parameter space (see Eq. (5) in Song *et al.* (2018b) for more details).

The range of intrinsic growth rates contained within the feasibility domain of a given community can be numerically approximated by the minimum and maximum values of such rates along the feasibility domain's border. To do so, for each species, we randomly selected 10000 intrinsic growth rates vectors in the border where the such species goes extinct, and then we extracted the the minimum and maximum values for each component of the $10000 \times S$ selected vectors, where $S$ is the number of species of the community. Intrinsic growth rates vectors where the $i$-th species goes extinct meet the following conditions: $\mathbf{r}_{\text{border i}} = -\mathbf{A}\mathbf{N}^*_{\text{border i}}$, where the components of the abundance vector (at a fixed point) are given by $N_i^* = 0$ and $N_j^* > 0$ for $j \neq i$. We explored this range for the feasibility domain of all our observed communities (Fig. S5). From this analysis we derived several insights, which agree with our theoretical expectations. First, the range of intrinsic growth rates allowed in the feasibility domain of communities of a single guild (left panels of Fig. S5) does not allow negative values. This is to be expected, since these communities only contain competitive interactions. As soon as inter-guild interactions are allowed (center and right panels), intrinsic growth rates can reach negative values and still be within the feasibility domain in some cases. Particularly, pollinator species seem to receive a larger net benefit from their inter-guild interactions, thus allowing a more negative minimum intrinsic growth rate than herbivores, on average. Likewise, maximum intrinsic growth rates allowed tend to be more constrained in communities with two or three guilds, so as not to generate overshoot dynamics from positive interactions feedbacks. As a last note, we also observe variation among the three parameterisations of intra-guild competition (the boxplots of different colors in Fig. S5). Again as expected, more structured interactions allow a wider range of intrinsic growth rates, which is consistent with these parameterisations displaying on average larger feasibility domains. Overall, we find that the range of intrinsic growth rates from the potentially feasible communities studied here align with ecological interpretations and with theoretical expectations. Note, however, that this exploration does not tell what are the specific growth rates of a given feasible community: in any realised community, intrinsic growth rates are not independent from each other, so they cannot be simply "sampled" from these ranges to form a feasible community. Rather, this brief analysis only delineates the limits below or above which feasibility is never observed.

#### Species exclusion ratio

To define the species exclusion ratio, first we define the probability of being the first species excluded from a feasible community when a large enough perturbation on the vector of intrinsic growth rates takes place. This metric, denoted as $P_i^E(\mathbf{A})$, quantifies the proportion of the feasibility domain's volume that is closer to the border where only a given species $i$ goes excluded ($N_i^* = 0$ and $N_j^* > 0$ for $i, j \in S$ and $j \neq i$). Its value for species $i$ can be calculated as:

$$P_i^E(\mathbf{A}) = \frac{\Omega\left(\mathbf{A}_i^M\right)}{\Omega\left(\mathbf{A}\right)}, \tag{S3}$$

where $\Omega\left(\mathbf{A}\right)$ represents the "size" (or relative volume) of the feasibility domain (given by Eqs. (5) and (6) in Song *et al.* (2018b)), $\mathfrak{I}$ denotes the *incenter* of the feasibility domain (on the unit-ball), and $\mathbf{A}_i^M$ is a modified interaction matrix, obtained by replacing the $i-$th column of $\mathbf{A}$ by the negative of the incenter's column vector. In a community with $S$ species, we define its incenter as

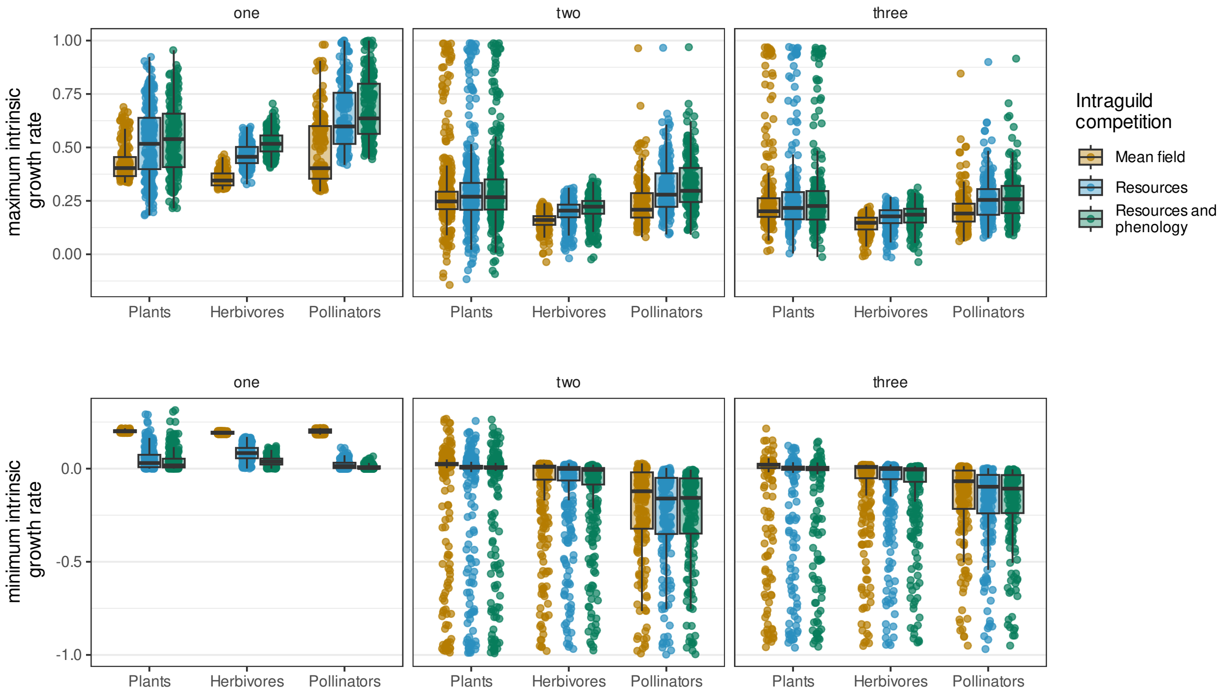


Fig. S5: Minimum values (lower panels) and maximum values (upper panels) of species intrinsic growth rates contained in the feasibility domains of each local community, aggregated by species guild and type of intra-guild competition. Horizontal panels show communities of one (left), two (center), and the full three guilds (right). As an example, take the communities with three guilds and the most structured intra-guild competition (resources and phenology): the range of intrinsic growth rates for plant species in these communities is the green boxplot of the right panels, with median values of $(0, 0.2)$.

the apex of the largest $S$-hyperspherical cap that can be placed in the intersection between the feasibility domain and the unit ball, and is tangent to each side of the feasibility domain's border. Then, this metric is weighted relative to a situation in which all species are equally likely to be the first species excluded:

$$ER_i(\mathbf{A}) = \frac{P_i^E(\mathbf{A})}{(1/S)}. \tag{S4}$$

The probability of extinction for single species in local communities is the complement of the probability of persistence defined in (Saavedra *et al.*, 2020), that is, $P(\text{persistence of } i) = 1 - P(\text{extinction of } i)$. In communities with $S$ species, it is possible to obtain analytically the probability of persistence by using the interaction matrix $\mathbf{A}$ and its $2^S - 2$ modifications (see (Saavedra *et al.*, 2020) for details). Nevertheless, for simplicity, $P(\text{persistence of } i)$ is usually estimated from numerical simulations of the Lotka–Volterra (LV) model, under several initial conditions, by calculating the proportion of fixed points (solutions) where the species $i$ abundance is positive, as in (Bartomeus *et al.*, 2021; Saavedra *et al.*, 2020). In our study, both approaches break due to the large number of species involved and the different interactions considered in our system. On the one hand, the 108 taxa identified in our study would require the evaluation of $2^{108} - 1 = 3.245186 \times 10^{32} - 1$ interaction matrices. If processing each matrix only took 1 mili-second, completing the calculation of $P(\text{persistence of } i)$ for all those species would require over $10^{16}$ million years. On the other hand, the combination of facilitative and competitive interactions hinders the convergence of LV dynamics to fixed points, because the predicted abundances of some species diverge and the extremely large numbers generated overflow the capability of ordinary differential equations solvers to handle them. Along with the above difficulties, it is worth mentioning that $P(\text{extinction of } i)$ is an absolute probability, meaning that it considers unfeasible scenarios, where part of the original community went extinct.

Even so, we wanted to make sure that the behavior of our $ER_i$ is qualitatively similar to $P(\text{extinction of } i)$ as defined in Saavedra *et al.* (2020). To do so, we have compared both metrics in networks that are easier to solve numerically. On the one hand, we used a set of 88 networks available in Mangal database (Poisot *et al.*, 2016) and computed their mean field interaction matrices, following the methodology and parameterisation described in Saavedra *et al.* (2016) for systems with no mutualistic trade-off. As can be seen in Fig. S6, the correlation between $ER_i$ and $P$(extinction of $i$) is excellent in that set of bipartite networks (Spearman $\rho = 0.965$, p-value $< 2.2 \times 10^{-16}$). On the other hand, we also tested that relation on the one-guild matrices present in our study system (see Fig. S6). Mean-field matrices show that $ER_i \simeq 1$ regardless of the probability of exclusion. This is expected because our mean-field matrices are circulant and, consequently, $P_i^E(\mathbf{A}) = 1/S$, where S is the number of species. Indeed, the observed values of $ER_i \neq 1$ are due to the small number of samples used to estimate $P_i^E(\mathbf{A})$ in Eq. (S3), via the quasi-Monte Carlo method proposed in (Song *et al.*, 2018b). Conceptually, an equal exclusion ratio for all species in a mean-field situation is a reassurance that in neutral situations our metric behaves adequately.

Regarding other one-guild interaction matrices, our results show a positive and significant correlation between the probabilities of exclusion and the exclusion ratios for floral visitors and herbivores (with $ER_i \lessapprox 1.6$), as in the case of Mangal networks. For plants, however, we observed a negative trend between both metrics, and the presence of larger exclusion ratios ($ER_i \in (0, 5)$). Indeed, the negative relation seems to be driven by species with $ER_i > 2$. According to our preeliminary analyses, the effect of other species on those with large $ER_i$ is very strong compared to their intraspecific competition. In that sense, the exclusion ratio we propose agrees with the expectation that stronger self-regulation implies a higher probability of persistence in a community. Further work is nevertheless needed to explore in more detail the limitation of the metric proposed here and its relationship with others like the one developed by Saavedra *et al.* (2020), but to summarise, the species exclusion ratio 1) can be reliably and efficiently estimated from complex communities with different interaction types, 2) shows homogeneity on the exclusion ratios of species in mean-field communities, as expected, and 3) agrees with theoretical expectations that larger inter-specific competition relative to self-regulation would increase the probability of exclusion.

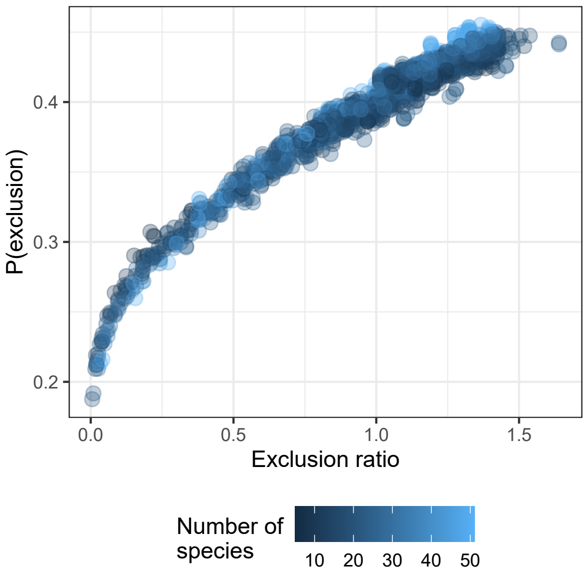


Fig. S6: Exclusion ratio and probability of exclusion from Saavedra *et al.* (2020) for a sample of 88 bipartite networks extracted from Mangal (Poisot *et al.*, 2016). To create the corresponding interaction matrices, we used the mean-field approach in (Saavedra *et al.*, 2016) and the following parameterisation: $\rho = 0.01$, $\delta = 0.00$ and $\gamma_0 = 0.50 \langle \gamma_{ij} \rangle$ (see further details on the parameters in the referred paper). Color of markers shows the number of species (or nodes) in each network.

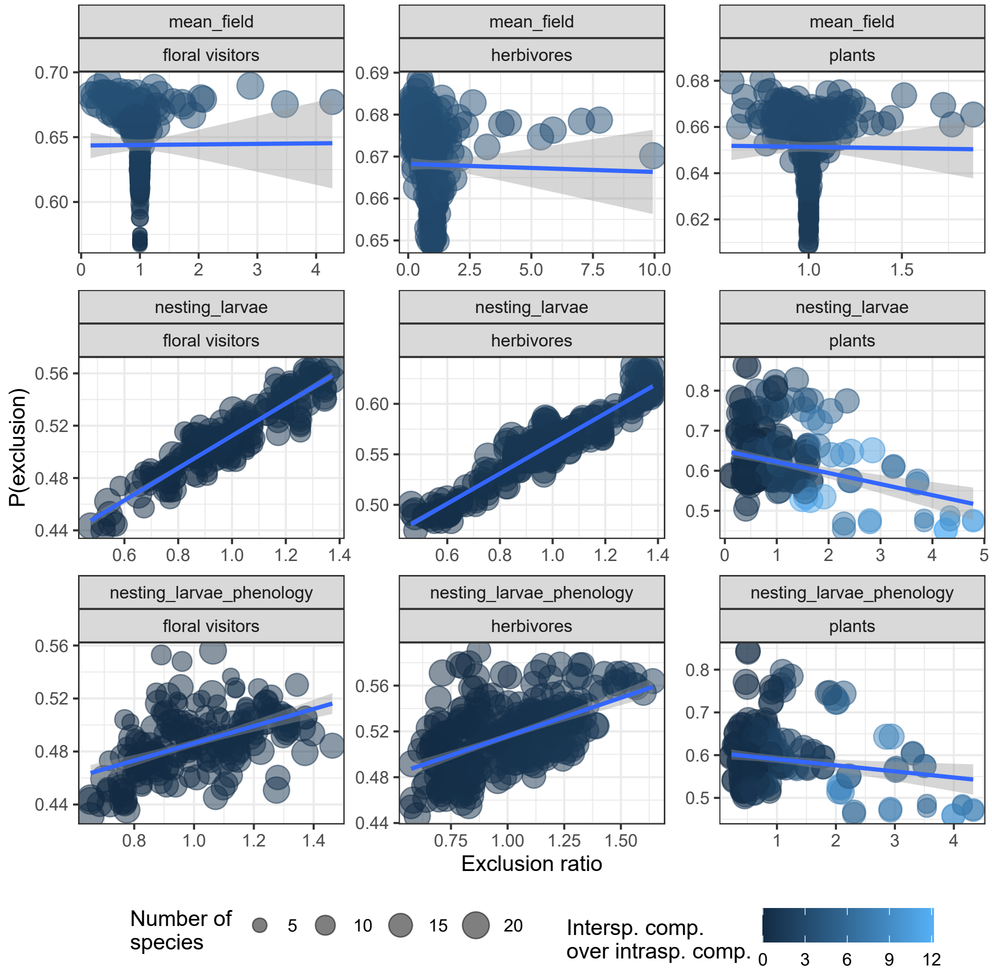


Fig. S7: Exclusion ratio and probability of exclusion from Saavedra *et al.* (2020) for those one-guild interaction matrices in our study system. Size of markers represents the number of species (or nodes) in each plot, whereas the color of markers shows the ratio between the sum of the effect of other species on the species represented by the marker and its intraspecific competition. Blue lines shows a linear regression between for the following model: P(exclusion of $i$) ˜ $ER_i$. Probabilities of exlcusion and exclusion ratios were estimated by using 10,000 directions of **r**-vectors sampled uniformly inside the full parameter space, respectively.

### Supplementary Section: Rarefaction analyses

We estimated the sampling coverage of plant-plant, plant-herbivore, and plant–pollinator interactions for each local community (combination of plot and year), pooling the data for all plant species, and using the coverage framework introduced in Chao *et al.* (2014). The sampling coverage of the observed interactions is simply their total relative abundances, or equivalently, the proportion of the total number of interactions in an assemblage that belong to interactions represented in the sample. To calculate the coverage in each plot, we used the R-package `iNEXT` v3.0 (Hsieh *et al.*, 2016). The average sampling coverage is $0.99 \pm 0.001$ for plant-plant interactions (Fig. S8), $0.97 \pm 0.02$ for plant-herbivore ones (Fig. S10), and $0.089 \pm 0.05$ for plant-pollinator ones (Fig. S9). This means that less than 5% of all interactions on average belong to undetected interactions. For this reason, extremely rare, undetected interactions do not make a significant contribution to that proportion, even if there are many such interactions. In addition, note that the above estimations do not account for the existence of forbidden interactions, that is, non-occurrences of pairwise interactions that can be accounted for by biological constraints, such as spatio-temporal uncoupling, size or reward mismatching, foraging constraints and physiological–biochemical constraints (Jordano, 2016).

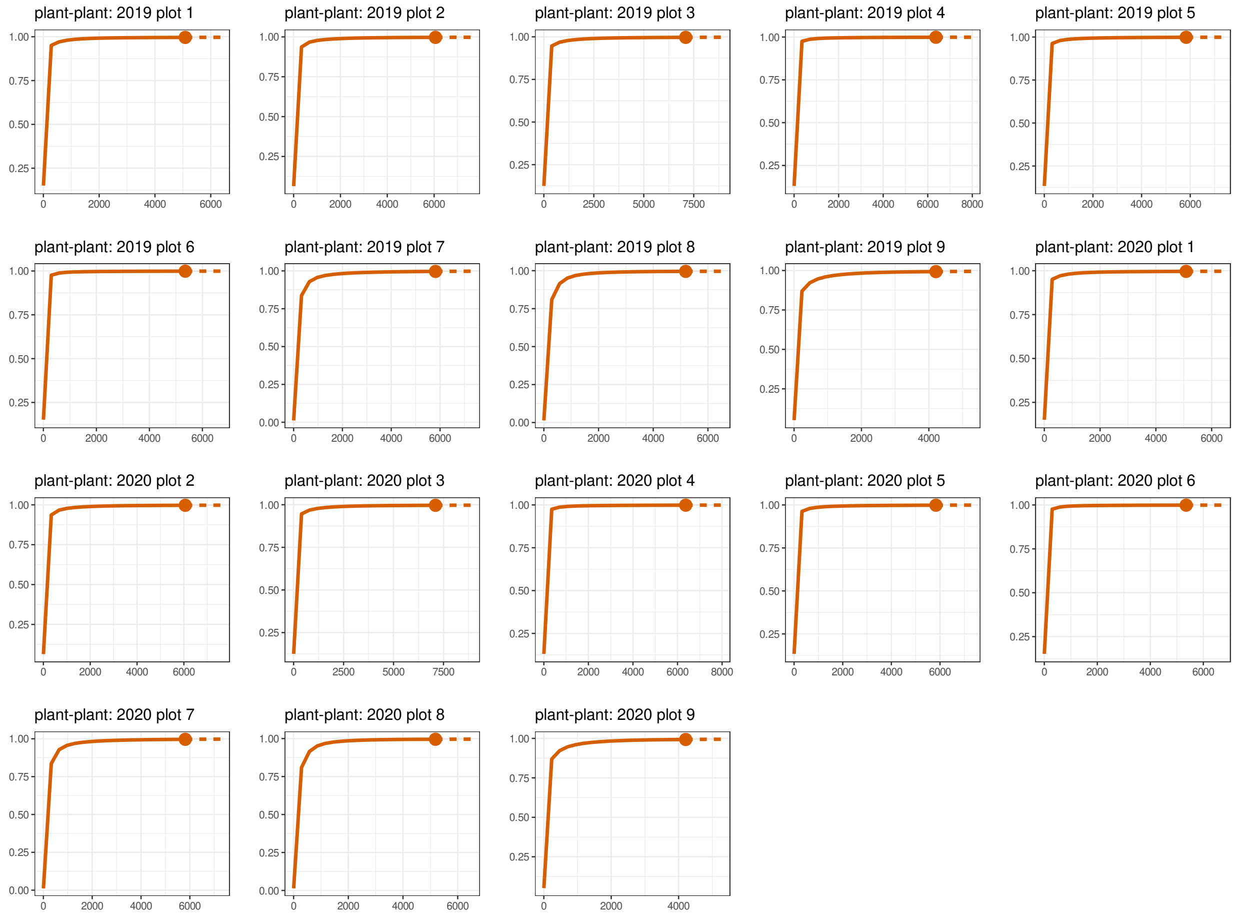


Fig. S8: Sample coverage for rarefied samples (solid line) and extrapolated samples (dashed line) as a function of sample size for plant-plant interactions sampled from each local community. The 95% confidence intervals were obtained by a bootstrap method based on 200 replications. Actual sampling coverage is denoted by solid dots.

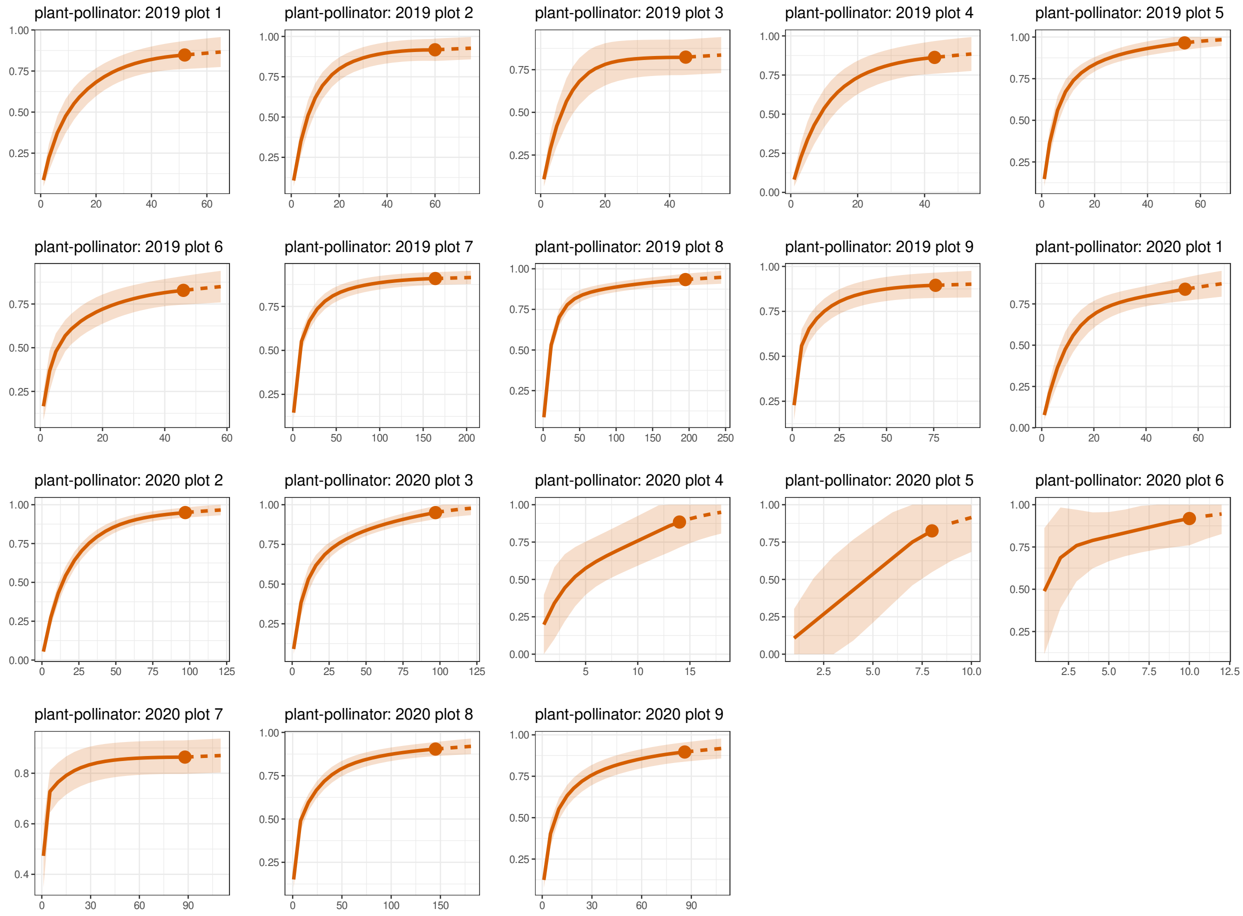


Fig. S9: Sample coverage for rarefied samples (solid line) and extrapolated samples (dashed line) as a function of sample size for plant-pollinator interactions sampled from each local community. The 95% confidence intervals were obtained by a bootstrap method based on 200 replications. Actual sampling coverage is denoted by solid dots.

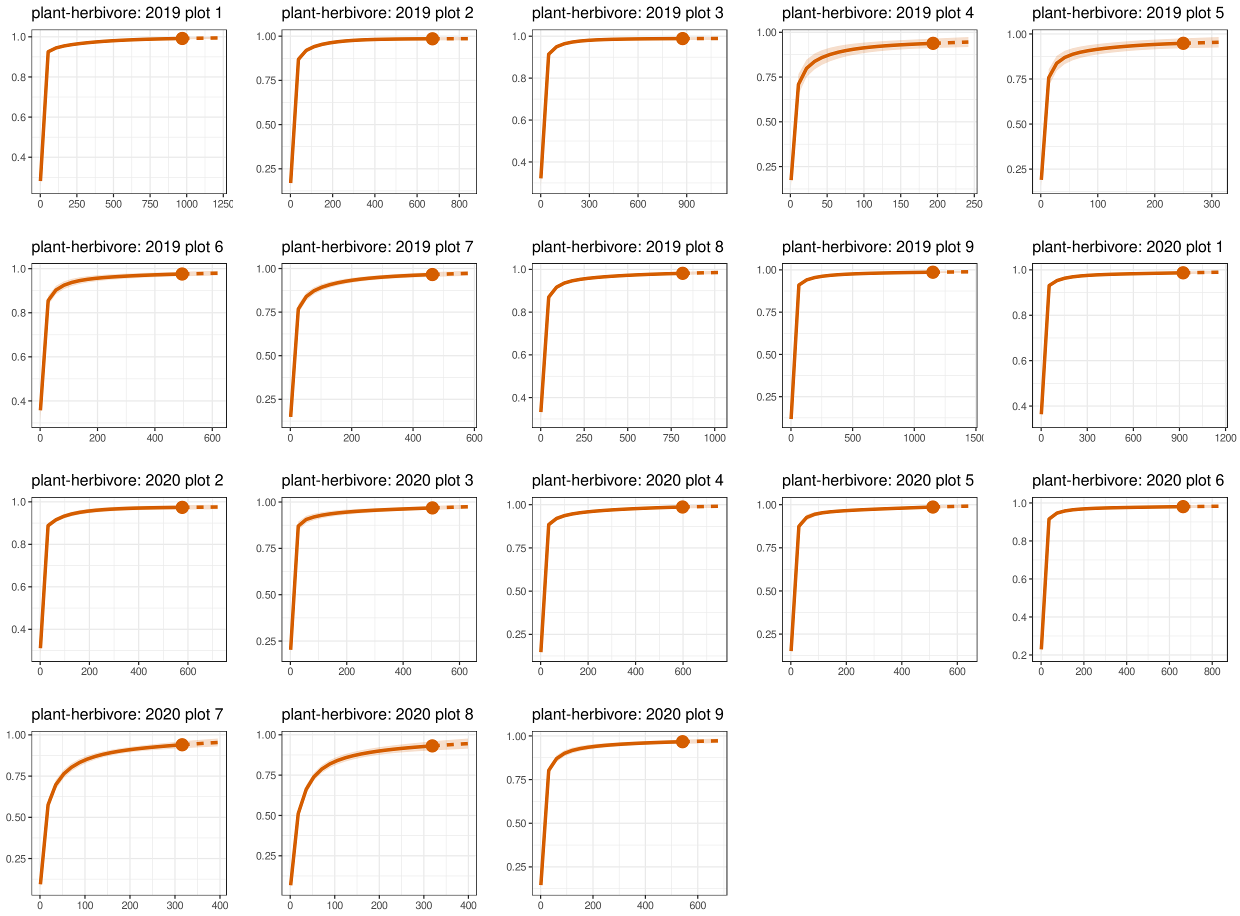


Fig. S10: Sample coverage for rarefied samples (solid line) and extrapolated samples (dashed line) as a function of sample size for plant-herbivore interactions sampled from each local community. The 95% confidence intervals were obtained by a bootstrap method based on 200 replications. Actual sampling coverage is denoted by solid dots.
